# Exportin-mediated nucleocytoplasmic transport maintains Pch2 homeostasis during meiosis

**DOI:** 10.1101/2023.07.10.548332

**Authors:** Esther Herruzo, Estefanía Sánchez-Díaz, Sara González-Arranz, Beatriz Santos, Jesús A. Carballo, Pedro A. San-Segundo

## Abstract

The meiotic recombination checkpoint reinforces the order of events during meiotic prophase I, ensuring the accurate distribution of chromosomes to the gametes. The AAA+ ATPase Pch2 remodels the Hop1 axial protein enabling adequate levels of Hop1-T318 phosphorylation to support the ensuing checkpoint response. While these events are focalized at chromosome axes, the checkpoint activating function of Pch2 relies on its cytoplasmic population. In contrast, forced nuclear accumulation of Pch2 leads to checkpoint inactivation. Here, we reveal the mechanism by which Pch2 travels from the cell nucleus to the cytoplasm to maintain Pch2 cellular homeostasis. Leptomycin B treatment provokes the nuclear accumulation of Pch2, indicating that its nucleocytoplasmic transport is mediated by the Crm1 exportin recognizing proteins containing Nuclear Export Signals (NESs). Consistently, leptomycin B leads to checkpoint inactivation and impaired Hop1 axial localization. Pch2 nucleocytoplasmic traffic is independent of its association with Zip1 and Orc1. We also identify a conserved functional NES in the non-catalytic N-terminal domain of Pch2 that is required for its nucleocytoplasmic traffic and proper checkpoint activity. In sum, we unveil another layer of control of Pch2 function during meiosis involving the nuclear export via the exportin pathway that is crucial to maintain the critical balance of Pch2 distribution among different cellular compartments.

## INTRODUCTION

In sexually-reproducing organisms, chromosome distribution to the gametes relies on a specialized type of cell division, called meiosis, which reduces the number of chromosomes to the half prior to fertilization (Bolcun-Filas & Handel, 2018). During the long meiotic prophase I, chromosomes undergo several elaborate and carefully-regulated processes: pairing, synapsis and recombination. Recombination involves the formation of developmentally programmed DNA double-strand breaks (DSBs) by the conserved Spo11 protein and its partners (Keeney *et al*, 2014). The interhomolog repair of a subset of these DSBs as crossovers (COs) is essential for accurate chromosome segregation (Hunter, 2015; Zickler & Kleckner, 2015; San-Segundo & Clemente-Blanco, 2020). This process requires chromosomes to undergo intimate interactions during meiotic prophase that are facilitated by active chromosome movements coordinated by cytoskeletal forces and the widely conserved LINC complex (Zeng *et al*, 2017; González-Arranz *et al*, 2020; Kim *et al*, 2022). Once each chromosome has paired with its homologous partner, a tripartite structure called the synaptonemal complex (SC) is assembled along the length of each chromosome pair to maintain a stable association between paired homologs. In *Saccharomyces cerevisiae*, Hop1, Red1 and Rec8 are structural components of the lateral elements of the SC (Hollingsworth *et al*, 1990; Smith & Roeder, 1997; Klein *et al*, 1999; Láscarez-Lagunas *et al*, 2020), and the transverse filament Zip1 protein composes the central region which holds together the axes (Sym *et al*, 1993). The central region also includes the central element made of the Ecm11 and Gmc2 proteins (Humphryes *et al*, 2013).

The fidelity of meiotic divisions is ensured by the meiotic recombination checkpoint, which blocks cell cycle progression at the end of prophase I while recombination intermediates (i.e., unrepaired DSBs) persist (Subramanian & Hochwagen, 2014). Briefly, the Mec1-Ddc2 sensor kinase is recruited to resected DSBs (Refolio *et al*, 2011) and phosphorylates Hop1 at various consensus S/T-Q sites in a Red1-dependent manner. Among the multiple S/T-Q sites in Hop1, phosphorylation of T318 (hereafter, Hop1-T318ph) is critical for checkpoint activation (Carballo *et al*, 2008; Lo *et al*, 2014; Penedos *et al*, 2015). The Pch2 AAA+ ATPase sustains adequate levels of Hop1-T318ph on chromosomes to signal checkpoint activity in *zip1*Δ (Herruzo *et al*, 2016). Hop1-T318ph, in turn, promotes the activation of Mek1 and its recruitment to chromosomes (Niu *et al*, 2005; Ontoso *et al*, 2013; Hollingsworth & Gaglione, 2019). Finally, the Mek1 effector kinase phosphorylates and inhibits the Ndt80 transcription factor leading to pachytene arrest (Chen *et al*, 2018). In addition, Mek1 acts on Rad54 and Hed1 helping to prevent intersister recombination (Niu *et al*, 2009; Callender *et al*, 2016).

Pch2 (known as TRIP13 in mammals) is a member of the AAA+ ATPase family; these AAA+ proteins use the energy provided by ATP hydrolysis to provoke conformational changes on their substrates (Hanson & Whiteheart, 2005; Puchades *et al*, 2020). The conserved Pch2 protein was initially identified in a screen for meiotic recombination checkpoint mutants in budding yeast (San-Segundo & Roeder, 1999), but in addition to the checkpoint it also participates in multiple meiotic processes regulating several aspects of CO recombination and chromosome morphogenesis both in yeast and other organisms (Bhalla & Dernburg, 2005; Li & Schimenti, 2007; Joshi *et al*, 2009; Joyce & McKim, 2009; Zanders & Alani, 2009; Farmer *et al*, 2012; Joshi *et al*, 2015; Lambing *et al*, 2015). Despite the varying effects resulting from the absence of Pch2 in different organisms, it has been proposed that the common meiotic function of Pch2 is the coordination of recombination with chromosome synapsis to ensure proper CO number and distribution (Bhalla, 2023).

The meiotic roles of Pch2 are exerted through its action on its preferred client: the HORMAD Protein Hop1 (Vader, 2015; Prince & Martinez-Perez, 2022). As a member of the HORMAD family, Hop1 contains a flexible safety belt in its HORMA domain, which enables it to adopt an open/unbuckled or closed conformation (Ye *et al*, 2017; West *et al*, 2018). The transition from closed-Hop1 to unbuckled-Hop1 is thought to be accomplished by Pch2 ATPase activity, poising Hop1 for binding to a closure motif in Hop1 itself, or in other proteins that interact with Hop1, like Red1 (Chen *et al*, 2014; Ye *et al*., 2017; Alfieri *et al*, 2018; Yang *et al*, 2020). In other organisms, Pch2^TRIP13^ requires the p31(COMET) adaptor, but no cofactor has yet been described for Pch2 action in yeast (Eytan *et al*, 2014; Ye *et al*, 2015; Balboni *et al*, 2020; Giacopazzi *et al*, 2020).

Pch2 localization studies feature a complicated scenario. Pch2 is highly enriched in the unsynapsed ribosomal DNA (rDNA) region. This rDNA-specific recruitment requires Orc1, which collaborates with Pch2 to exclude Hop1 from the nucleolus thus limiting meiotic DSB formation at the repetitive rDNA array (San-Segundo & Roeder, 1999; Vader *et al*, 2011). Pch2 is also detected as individual foci colocalizing with Zip1 on synapsed chromosomes (San-Segundo & Roeder, 1999; Joshi *et al*., 2009). Targeting of Pch2 to the SC depends on Zip1, but other factors such as RNAPII-dependent transcription (Cardoso da Silva *et al*, 2020), Top2 (Heldrich *et al*, 2020), Nup2 (Subramanian *et al*, 2019) or chromatin modifications driven by Sir2 and Dot1 (San-Segundo & Roeder, 2000; Ontoso *et al*., 2013; Cavero *et al*, 2016), also influence Pch2 chromosomal distribution. Pch2 recruitment to chromosomes removes Hop1 from the axes (Borner *et al*, 2008; Herruzo *et al*., 2016; Subramanian *et al*, 2016), likely by disrupting Hop1-Red1 interaction via its remodeling activity towards the HORMA domain (West *et al*., 2018). Hop1 removal downregulates DSB formation on chromosomes that have successfully identified their homologs and formed COs, and silences the meiotic recombination checkpoint (Raina & Vader, 2020; Herruzo *et al*, 2021). In addition to the nucleolar and chromosomal population, a cytoplasmic supply of Pch2 also exists; this cytoplasmic pool is necessary and sufficient to sustain meiotic checkpoint activation (Herruzo *et al*, 2019; Herruzo *et al*., 2021). Based on all these observations, and the demonstrated role of Pch2 in promoting Hop1 chromosomal incorporation and T318 phosphorylation in *zip1Δ* (Herruzo *et al*., 2016), a model emerges in which Pch2, from the cytoplasm, structurally remodels Hop1 by using its ATPase activity, ensuring that unbuckled Hop1 is available to associate with Red1 via the closure motif determining its axial incorporation and achieving proper phosphorylation levels at T318 to support checkpoint activity (Herruzo *et al*., 2021).

The importance of a precise localization of Pch2 for an accurate meiotic checkpoint response argues that its distribution between the different cellular compartments must be finely regulated. Here, we combine detailed cytological, molecular and genetics studies to provide novel insights into how Pch2 travels from the nucleus to the cytoplasm, thereby contributing to delineate the regulatory network governing Pch2 subcellular distribution and function.

## RESULTS and DISCUSSION

### Pch2 nuclear export is mediated by the Crm1 exportin

The existence of at least three different subpopulations of Pch2 residing in the nucleolus, chromosomes and cytoplasm, together with the recent observation that an exquisite balance of Pch2 subcellular distribution is crucial to maintain a proper meiotic recombination checkpoint response (Herruzo *et al*., 2021), suggest that the nucleocytoplasmic transport of Pch2 must be tightly controlled. As an initial approach to explore this mechanism, we analyzed if Pch2 travels from the nucleus to the cytoplasm via Crm1, which is the main exportin mediating the nuclear export of proteins containing a Nuclear Export Signal (NES) (Stade *et al*, 1997). The drug leptomycin B (LMB) is a powerful tool to establish whether the nuclear export of a NES containing protein is mediated by the CRM1/XPO1/KAP124 exportin pathway. LMB binds to CRM1 disrupting the formation of the trimeric NES-CRM1-RanGTP export complex required for the transport from the nucleus to the cytoplasm (Sun *et al*, 2013). Thus, LMB treatment leads to the nuclear accumulation of NES-containing proteins. In *Schizosaccharomyces pombe* and mammalian cells, LMB is capable of binding directly to CRM1 inhibiting nuclear export, but *S. cerevisiae* cells are resistant to LMB because the wild-type budding yeast Crm1 does not bind to the drug. However, the Crm1-T539C mutant version is capable of binding LMB with high affinity and renders *S. cerevisiae* cells sensitive to LMB (Neville & Rosbash, 1999). Thus, to determine whether Pch2 travels from the nucleus to the cytoplasm via the Crm1 pathway, we analyzed Pch2 subcellular localization after LMB treatment in a *crm1-T539C* mutant background (Figure 1A-D). First, we confirmed that control *crm1-T539C* diploid cells, in the absence of LMB, completed meiosis and sporulation with fairly normal kinetics and efficiency (Figure S1A, S1B); however, addition of LMB during prophase I (15h) blocked meiotic progression (Figure S1B), indicating that, as expected, nuclear export is required for completion of meiosis and sporulation. More important, we also checked that the untreated *zip1Δ crm1-T539C* mutant showed a strong meiotic arrest (Figure S1A, S1B), indicating that the *crm1-T539C* mutation itself, in the absence of LMB, does not alter the *zip1Δ*-induced checkpoint response. Thus, the *crm1-T539C* mutant is a valid tool to explore the implication of nuclear export in Pch2 subcellular distribution in *S. cerevisiae*.

**Figure 1.**
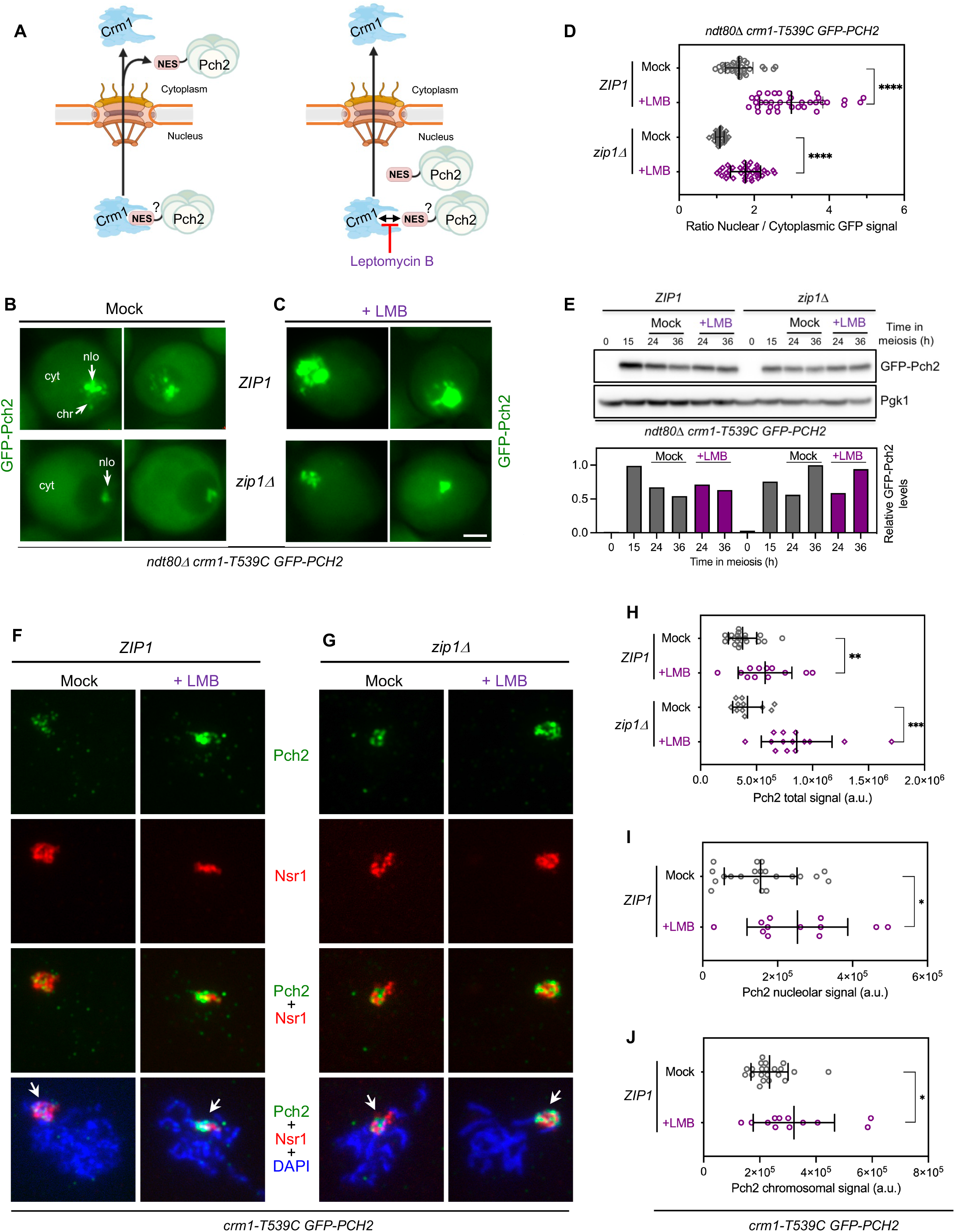
Nuclear export of Pch2 is blocked by leptomycin B (LMB). **(A)** Schematic representation of protein nuclear export via nuclear pore complexes mediated by the interaction of a Nuclear Export Sequence (NES) with the Crm1 exportin (see text for details). LMB blocks NES binding to Crm1 preventing nuclear export. The presence of a putative NES in Pch2 is depicted. For simplicity, the NES is drawn only in one subunit of the Pch2 hexamer. BioRender.com was used to create this figure. **(B, C)** Fluorescence microscopy images of GFP-Pch2 localization in *ZIP1* and *zip1Δ* cells, mock-treated (B), or treated with 500 ng/ml LMB (C), 15 h after meiotic induction. Images were taken at 24 h. Two representative individual cells for each condition are shown. Localization of GFP-Pch2 in the nucleolus (nlo), chromosomes (chr), and cytoplasm (cyt) is indicated. Scale bar, 2 μm. Strains are: DP1927 (*ndt80Δ crm1-T539C GFP-PCH2*) and DP1837 (*zip1Δ ndt80Δ crm1-T539C GFP-PCH2*). **(D)** Quantification of the ratio of nuclear (including nucleolar) to cytoplasmic GFP fluorescent signal for the experiment shown in (B, C). Error bars, SD. **(E)** Western blot analysis of GFP-Pch2 production (using anti-Pch2 antibodies) in the absence (mock) or presence of LMB. Pgk1 was used as loading control. The graph shows the quantification of GFP-Pch2 relative levels. **(F, G)** Immunofluorescence of spread meiotic chromosomes at pachytene stained with anti-Pch2 antibodies (to detect GFP-Pch2; green), anti-Nsr1 antibodies (red) and DAPI (blue). Representative *ZIP1* (F) and *zip1Δ* (G) nuclei, mock-treated, or treated with 500 ng/ml LMB 15 h after meiotic induction are shown. Arrows point to the rDNA region. Spreads were prepared at 19 h. Scale bar, 2 μm. Strains are: DP1717 (*crm1-T539C GFP-PCH2*) and DP1721 (*zip1Δ crm1-T539C GFP-PCH2*). **(H)** Quantification of the GFP-Pch2 total signal for the experiment shown in (F, G). Error bars, SD; a.u., arbitrary units. **(I-J)** Quantification of the nucleolar (I) and chromosomal (J) GFP-Pch2 signal for the experiment shown in (F). Error bars, SD; a.u., arbitrary units.

Pch2 localization was assessed by fluorescence microscopy analysis of a functional version of GFP-tagged Pch2 expressed from the previously described *P_HOP1_-GFP-PCH2* construct (Herruzo *et al*., 2019; Herruzo *et al*., 2021), referred to as *GFP-PCH2* throughout the article for simplicity. To avoid side effects resulting from differences in meiotic progression, GFP-Pch2 subcellular distribution was studied in prophase I-arrested *ndt80Δ crm1-T539C* live meiotic cells in otherwise wild-type (*ZIP1)* and *zip1Δ* backgrounds. Cells were treated with LMB, or mock-treated with the solvent ethanol, as control. As previously described, in mock treated wild-type *ZIP1* cells, Pch2 localized to one side of the nucleus in a region corresponding to the nucleolus, it was also detected as discrete faint nuclear foci corresponding to the synapsed chromosomes, and it showed diffuse homogenous cytoplasmic signal as well (Figure 1B, top panels). Addition of LMB led to a strong accumulation of Pch2 in the nucleus displaying a much more intense signal especially in the presumed nucleolus, but also on the chromosomes and even in the nucleoplasm (Figure 1C, top panels). On the other hand, in the *zip1Δ* untreated control, Pch2 was only present in the nucleolus and the cytoplasm (Figure 1B, bottom panels), but LMB addition resulted in a prominent nucleolar accumulation and nucleoplasmic distribution, concomitant with reduced cytoplasmic localization (Figure 1C, bottom panels). Accordingly, the ratio between nuclear (including nucleolus) and cytoplasmic GFP signal increased after LMB treatment both in *ZIP1* or *zip1Δ* strains (Figure 1D). Despite the altered subcellular distribution, total GFP-Pch2 protein levels remained unchanged in the presence or absence of LMB (Figure 1E).

To obtain more detailed information, we also analyzed the localization of GFP-Pch2 by immunofluorescence on pachytene chromosome spreads after LMB treatment. We first describe the distribution pattern in *ZIP1* strains. In the absence of LMB, and consistent with the well-characterized localization of Pch2 (San-Segundo & Roeder, 1999; Joshi *et al*., 2009; Herruzo *et al*., 2016) and the observations in live meiotic cells (Figure 1B), the GFP-Pch2 protein localized mainly in the rDNA region, marked by the nucleolar protein Nsr1 (Figure 1F). The chromosome-associated population of Pch2, which is only present in synapsis-proficient strains, was not easily detectable with this technique in the BR strain background, as previously reported (Herruzo *et al*., 2016; Herruzo *et al*., 2019) (Figure 1F). However, the addition of LMB led to a general increase of GFP-Pch2 amount on the spread nuclei (Figure 1H), affecting both the nucleolar (Figure 1I) and the chromosomal (Figure 1J) populations. Moreover, in LMB treated *ZIP1* cells, the accumulation of Pch2 in the nucleus was also associated to the increased formation of polycomplexes, which are organized extrachromosomal assemblies of SC components (Dong & Roeder, 2000), in 29.4% of nuclei (Figure S2A, S2C). On the other hand, on spread chromosomes of the *zip1Δ* mutant, Pch2 is only associated to the rDNA region; the addition of LMB led to an increased GFP-Pch2 signal restricted to the nucleolar area (Figure 1G, 1H).

In sum, these results demonstrate that inhibition of the Crm1 exportin pathway leads to Pch2 nuclear accumulation indicating that Pch2 travels from the nucleus to the cytoplasm using this pathway.

### Effect of Crm1-mediated export inhibition on checkpoint function

We have previously described that the main role of Pch2 in the *zip1Δ-*induced checkpoint is to promote Hop1 association to unsynapsed chromosome axes supporting high levels of Hop1 phosphorylation at T318, which in turn sustain Mek1 activation (Herruzo *et al*., 2016). Artificial redirection of Pch2 to different subcellular compartments (i.e., forced nuclear accumulation) impairs these functions (Herruzo *et al*., 2021). To determine whether inhibition of the Crm1 nuclear export pathway, which alters Pch2 subcellular distribution, affects checkpoint function in *zip1Δ*, we examined the impact on Hop1 incorporation onto chromosome axes as well as the checkpoint status using Hop1-T318 phosphorylation as marker for Pch2-dependent checkpoint activity (Carballo *et al*., 2008; Penedos *et al*., 2015; Herruzo *et al*., 2016).

Analysis of Hop1 localization by immunofluorescence of *zip1Δ* spread nuclei in the absence of LMB revealed its characteristic intense and continuous signal along the unsynapsed axes, and its exclusion from the rDNA region containing Pch2 (Figure 2A). In contrast, after LMB treatment, Hop1 localization was compromised, displaying reduced intensity and a less continuous axial pattern (Figure 2A, 2B). To elude the possible effect of the different kinetics of meiotic progression, emphasized by the meiotic block conferred by the addition of LMB (Figure S1), we used prophase I-arrested *ndt80Δ* strains for an accurate analysis of Hop1-T318 phosphorylation levels without interference from cell cycle progression. As observed in Figure 2C, in the absence of LMB, the checkpoint was active in *zip1Δ* cells, as manifested by the high levels of Hop1-T318ph, compared to the wild type (*ZIP1)*. However, phosphorylation levels of this checkpoint marker were notably reduced by the addition of LMB in *zip1Δ* cells (Figure 2C).

**Figure 2.**
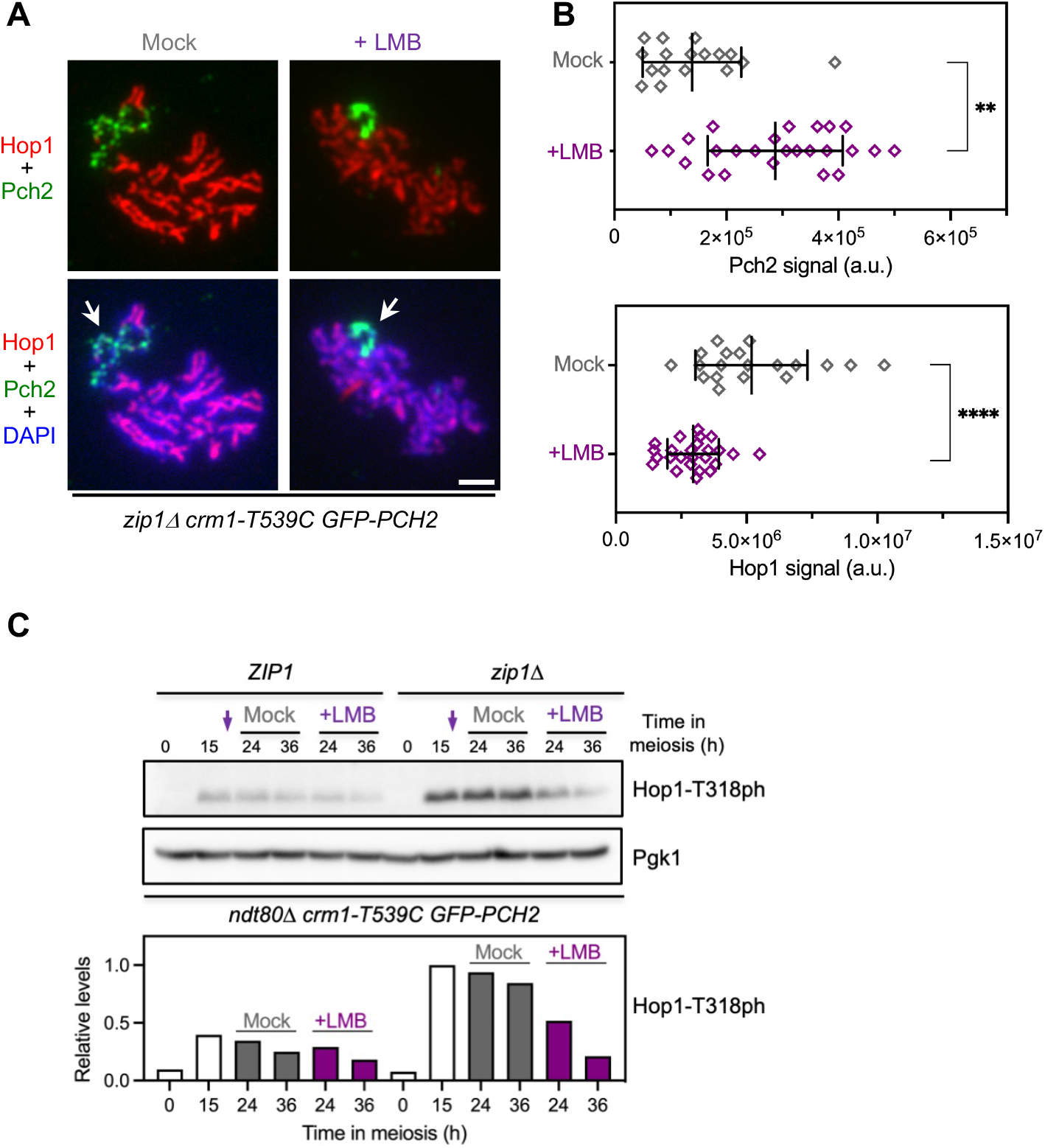
LMB treatment impairs the meiotic recombination checkpoint. **(A)** Immunofluorescence of *zip1Δ* spread meiotic chromosomes stained with anti-Hop1 (red) and anti-GFP (to detect GFP-Pch2; green) antibodies. Representative nuclei, either mock-treated, or treated with 500 ng/ml LMB 15 h after meiotic induction, are shown. Arrows point to the rDNA region. Spreads were prepared at 19 h. Scale bar, 2 μm. The strain is DP1721 (*zip1Δ crm1-T539C GFP-PCH2*). **(B)** Quantification of Pch2 (top graph) and Hop1 (bottom graph) signal for the experiment shown in (A). Error bars, SD; a.u., arbitrary units. **(C)** Western blot analysis of Hop1-T318 phosphorylation (ph) in the absence (mock) or presence of LMB added at 15 h after meiotic induction (arrows). Pgk1 was used as loading control. The graph shows the quantification of Hop1-T318ph relative levels. Strains are: DP1927 (*ndt80Δ crm1-T539C GFP-PCH2*) and DP1837 (*zip1Δ ndt80Δ crm1-T539C GFP-PCH2*).

Thus, the accumulation of Pch2 in the nucleus of *zip1Δ* cells imposed by inhibition of Crm1 correlates with defects in Hop1 localization and phosphorylation, and inactivation of the meiotic recombination checkpoint. We cannot rule out that the nuclear accumulation of other factors involved in the checkpoint besides Pch2 could be responsible for the defective response observed after LMB treatment. For example, nucleocytoplasmic transport also appears to be involved in Ndt80 regulation (Wang *et al*, 2011). Nonetheless, we point out that the forced nuclear accumulation of Pch2 alone by fusion to a strong NLS leads to checkpoint downregulation (Herruzo *et al*., 2021), arguing that the simply block of Pch2 nuclear export by itself may account for the impaired checkpoint activity observed in LMB-treated *zip1Δ* cells.

### Pch2 nucleocytoplasmic traffic is independent of Zip1 and Orc1

Since Orc1 is required for Pch2 nucleolar targeting (Vader *et al*., 2011), to further delineate the requirements for Pch2 nucleocytoplasmic transport we analyzed the effect of depleting Orc1 under LMB treatment conditions. That is, we explored if Pch2 nucleocytoplasmic traffic involves its transit through the rDNA region. To this end, we used the *orc1-3mAID* degron allele previously described; in this mutant, Pch2 does not localize to the nucleolus because of Orc1 degradation induced by auxin (Herruzo *et al*., 2019). We studied the localization of GFP-Pch2 in *orc1-3mAID* meiotic live cells and on chromosome spreads of both *ZIP1* and *zip1Δ* strains. All the experiments involving *orc1-3mAID* were performed with auxin added at 12 h after meiotic induction to induce Orc1 depletion during prophase I. For clarity, we first describe the localization patterns in *ZIP1* cells. In most mock-treated cells of the *orc1-3mAID* mutant, GFP-Pch2 was only detected in the cytoplasm and on nuclear foci or lines of different sizes likely corresponding to the chromosomes (Figure 3A, mock panel; Figure S2D). Indeed, immunofluorescence analyses of nuclear spreads confirmed the lack of Pch2 in the nucleolus (marked by Nsr1) and its presence on the chromosomes displaying a dotty-linear signal in the LMB-untreated control (Figure 3C, mock panels). In contrast, in the majority of LMB-treated *ZIP1* cells, the cytoplasmic signal was reduced and GFP-Pch2 displayed a robust nuclear localization with accumulation in a strong large focus that could be combined with a dotty or linear pattern (Figure 3A, 3B; Figure S2D). In a small fraction of cells (16-20%), a diffuse nuclear signal could also be found in the presence or absence of LMB. This minor pattern appeared to be exclusive of *orc1-3mAID crm1-T539C* cells (Figure S2D). Analyses of nuclear spreads revealed that, in the presence of LMB, the GFP-Pch2 signal was much more intense, showing a more continuous linear chromosomal pattern (Figure 3C, left LMB panel; Figure 3E). Moreover, 45% of the spread nuclei showed a very intense Pch2 focus that did not correspond to the nucleolus (Figure 3C, right LMB panels), but colocalized with Zip1 at polycomplexes (Figure S2B, S2C). In these cases, the chromosomal staining of Zip1 and Pch2 was somewhat masked by the strong signal of the polycomplex. Thus, we conclude from these results that the block of nucleocytoplasmic traffic in Orc1-depleted cells, lacking nucleolar Pch2, leads to a stronger association of Pch2 with the SC or assemblies of SC components. Next, to determine whether the nuclear retention of Pch2 upon LMB treatment requires its interaction with Zip1, we analyzed Pch2 localization in the *zip1Δ orc1-3mAID* mutant. We observed that GFP-Pch2 localized exclusively in the cytoplasm in the absence of LMB (Figure 3A, mock panel) but, upon LMB treatment, Pch2 was retained in the nucleus displaying a diffuse nucleoplasmic distribution (Figure 3A, LMB panel, Figure 3B). There was no chromatin-associated Pch2 signal in either untreated or LMB-treated nuclei (Figure 3D, 3E). In sum, these observations suggest that shuttling of Pch2 between the nucleus and the cytoplasm involves neither the SC (Zip1-dependent) nor the nucleolar (Orc1-dependent) association of Pch2.

**Figure 3.**
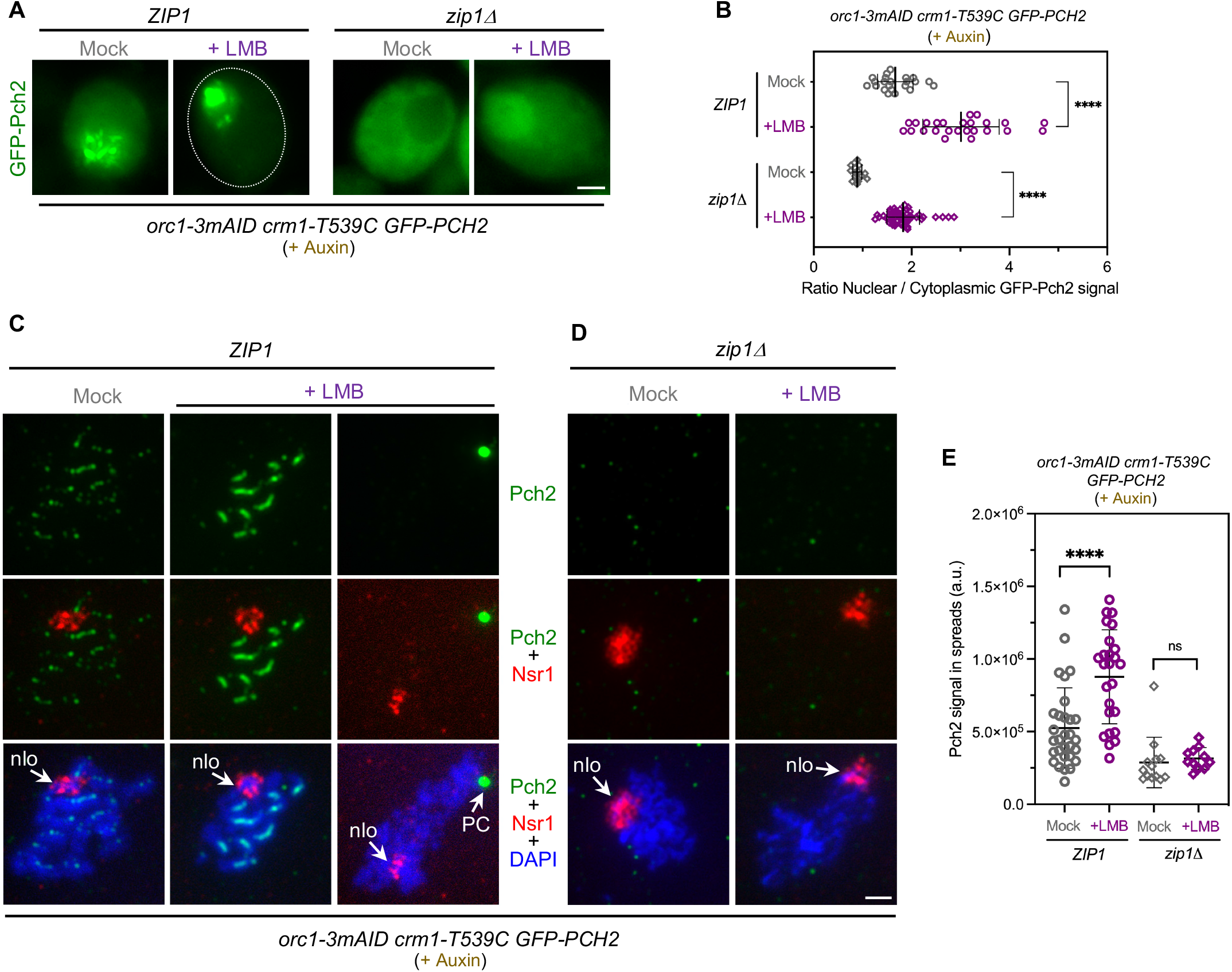
Pch2 nucleocytoplasmic traffic does not involve its association with Orc1 and Zip1. **(A)** Fluorescence microscopy images of GFP-Pch2 localization in *ZIP1 orc1-3mAID* and *zip1Δ orc1-3mAID* cells, mock-treated, or treated with 500 ng/ml LMB, 15 h after meiotic induction. Auxin (500μM) was added to the cultures 12 h after meiotic induction to induce Orc1 degradation. Images were taken at 19 h. Cells representing the predominant GFP-Pch2 localization pattern for each condition are shown. Additional examples of GFP-Pch2 distribution are presented in Figure S2D. Scale bar, 2 μm. Strains are: DP1885 (*orc1-3mAID crm1-T539C GFP-PCH2*) and DP1886 (*zip1Δ orc1-3mAID crm1-T539C GFP-PCH2*). **(B)** Quantification of the ratio of nuclear (including nucleolar) to cytoplasmic GFP fluorescent signal for the experiment shown in (A). Error bars, SD. **(C-D)** Immunofluorescence of *ZIP1 orc1-3mAID* (C) and *zip1Δ orc1-3mAID* (D) spread meiotic chromosomes stained with anti-Pch2 (green) and anti-Nsr1 (red) antibodies, and DAPI (blue). Representative nuclei, either mock-treated, or treated with 500 ng/ml LMB 15 h after meiotic induction, are shown. Auxin (500 μM) was added to the cultures 12 h after meiotic induction to induce Orc1 degradation. Spreads were prepared at 19 h. Arrows point to the nucleolar region (nlo) and the Polycomplex (PC). Scale bar, 2 μm. The strains are the same used in (A). **(D)** Quantification of Pch2 signal for the experiment shown in (C-D). Error bars, SD; a.u., arbitrary units.

### A NES sequence in the NTD of Pch2 drives its export out of the nucleus

It is well known that the Crm1 exportin binds proteins possessing Nuclear Export Signals (NESs) for its transport out of the nucleus (Fung *et al*, 2021). We have observed that Pch2 export depends on Crm1 but, in principle, Pch2 could travel bound to putative partner(s) containing NESs or via NES(s) present in Pch2 itself. Thus, we searched for consensus NESs in the Pch2 sequence using the LocNES prediction tool (Xu *et al*, 2015) combined with visual matching to known patterns for hydrophobic residues distribution in NESs (Fung *et al*, 2017). Three putative NESs with high score probability were predicted; all of them present in the N-terminal domain (NTD) of Pch2 at amino acid positions 98-107, 127-136 and 205-214. To determine whether these putative NESs are functionally relevant for Pch2 nucleocytoplasmic traffic, we generated three GFP-tagged Pch2 versions harboring substitutions of hydrophobic residues (Leu, Ile, Val or Phe) present in each predicted putative NES to alanine (*pch2-ntd^98-107^-6A*, *pch2-ntd^127-136^-5A* and *pch2-ntd^205-214^-4A*) (Figure S3A). Centromeric plasmids containing these constructs were transformed into a *zip1Δ pch2Δ* strain and the localization of GFP-Pch2 was examined in live meiotic cells. The *pch2-ntd^98-107^-6A* and *pch2-ntd^127-136^-5A* mutations both abolished nucleolar localization and led to an exclusively cytoplasmic distribution of Pch2 (Figure S3A, S3B). These mutations lie in the extended region of the NTD required for Orc1 interaction (Villar-Fernández *et al*, 2020), thus explaining the defective nucleolar targeting; therefore, they were discarded for further analyses. However, the GFP-pch2-ntd^205-214^-4A version showed a strong accumulation inside the nucleus as expected if a functional NES is disrupted (Figure S3A, S3B). Therefore, we generated strains harboring the *GFP-pch2-ntd^205-214^-4A* mutation (hereafter named as *pch2-nes4A*) integrated at the *PCH2* genomic locus for more detailed analyses. We confirmed the significant increase in nuclear localization both in the nucleolus and nucleoplasm of the Pch2-nes4A version (Figure 4A, 4B). Consistent with the fact that Pch2 nuclear buildup in *zip1Δ* cells is deleterious for the checkpoint (Herruzo *et al*., 2021), the *GFP-pch2-nes4A* mutant suppressed the meiotic block of *zip1Δ* though to a lesser extent than the *zip1Δ pch2Δ* mutant did (Figure 4C). Furthermore, the levels of Hop1-T318 phosphorylation were reduced in the *zip1Δ GFP-pch2-nes4A* double mutant compared to *zip1Δ* (Figure 4D) reflecting impaired checkpoint activity. To confirm that these checkpoint defects stem from the impaired Pch2 nucleocytoplasmic transport resulting from a defective NES and not from other possible alterations in Pch2 activity and/or structure, we added an ectopic bona-fide strong NES from the PKI protein (Wen *et al*, 1995) generating a *GFP-NES^PKI^-pch2-nes4A* version. Importantly, the *GFP-NES^PKI^-pch2-nes4A* construct alleviated the nuclear accumulation of GFP-Pch2-nes4A (Figure 4A, 4B) and reestablished checkpoint function, as revealed by the restoration of the meiotic block (Figure 4C) and higher Hop1-T318 phosphorylation levels especially at the later time point (48 h) (Figure 4D). Thus, we conclude that the ^205^**L**STE**F**DK**I**D**L**^214^ sequence in the Pch2 NTD domain (designated as NES^Pch2^) behaves as a functional NES matching the Class 1a signature (Fung *et al*., 2017). We demonstrate that NES^Pch2^ drives the nuclear export of Pch2, and it is critical for maintaining an exquisite balance of Pch2 subcellular distribution essential for a precise meiotic recombination checkpoint response. These findings also indicate that Pch2 does not need additional partners, besides the nuclear export machinery, to travel from the nucleus to the cytoplasm where it exerts its essential checkpoint activating function.

**Figure 4.**
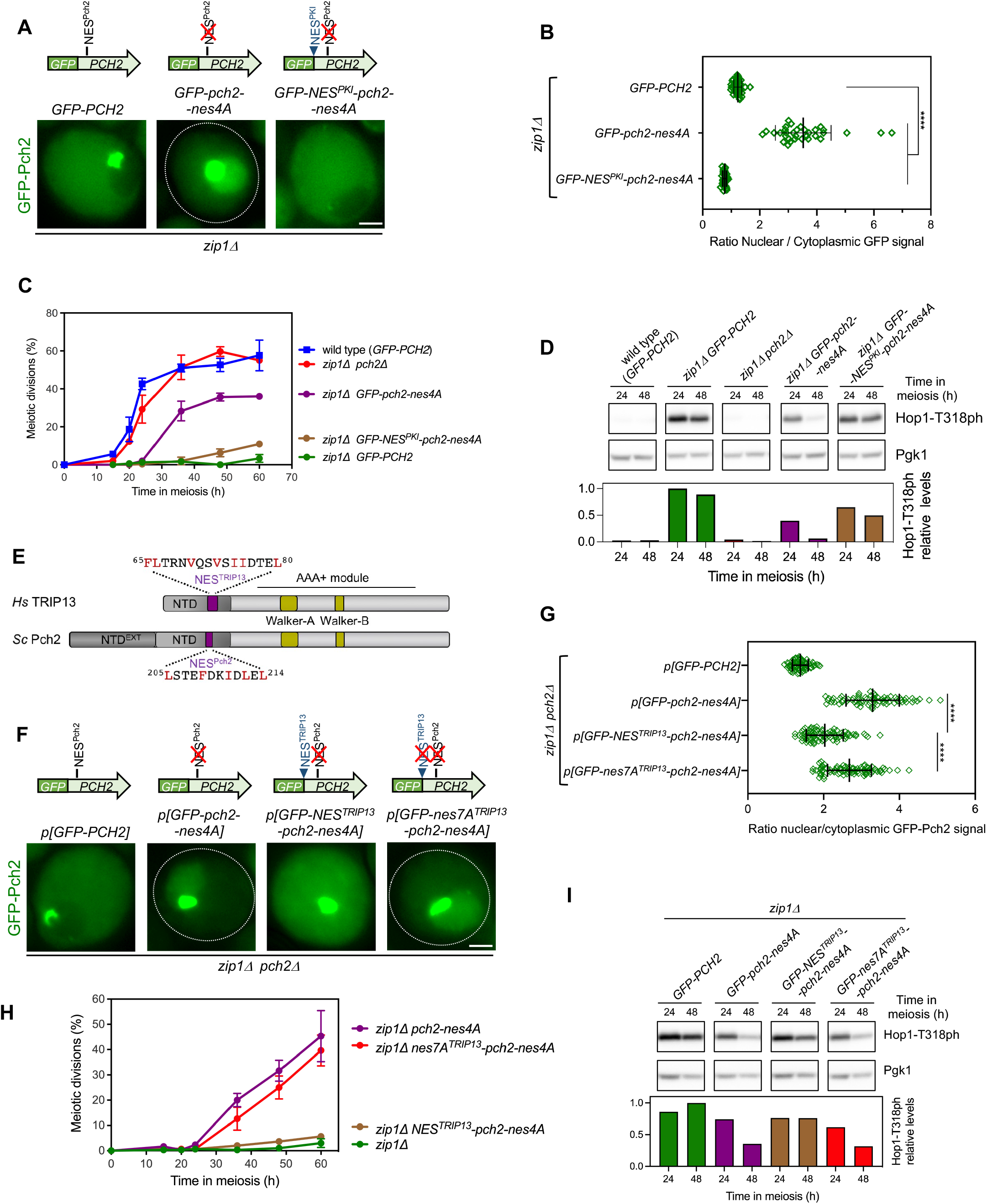
A conserved functional NES in the NTD of Pch2 drives its nuclear export. **(A)** Fluorescence microscopy images of the localization of GFP-Pch2, GFP-Pch2-nes4 and GFP-NES^PKI^-Pch2-nes4A in *zip1Δ* cells. Images were taken 15 h after meiotic induction. Representative cells are shown. A schematic representation of the different GFP-Pch2 versions is also shown on top. Scale bar, 2 μm. Strains are: DP1625 (*zip1Δ GFP-PCH2*), DP1986 (*zip1Δ GFP-pch2-nes4*) and DP1992 (*zip1Δ GFP-NES^PKI^-pch2-nes4A*). **(B)** Quantification of the ratio of nuclear (including nucleolar) to cytoplasmic GFP fluorescent signal for the experiment shown in (A). Error bars, SD. **(C)** Time course analysis of meiotic nuclear divisions; the percentage of cells containing two or more nuclei is represented. Error bars: SD; n=3. At least 300 cells were scored for each strain at every time point. Strains are: DP1624 (*GFP-PCH2*), DP1625 (*zip1Δ GFP-PCH2*), DP1029 (*zip1Δ pch2Δ*), DP1986 (*zip1Δ GFP-pch2-nes4A*) and DP1992 (*zip1Δ GFP-NES^PKI^-pch2-nes4A*). **(D)** Western blot analysis of Hop1-T318 phosphorylation in the strains analyzed in (C) at the indicated time points in meiosis. Pgk1 was used as a loading control. The graph shows the quantification of Hop1-T318ph relative levels. **(E)** Schematic representation of the *S. cerevisiae* Pch2 protein (ScPch2) and the and human ortholog (HsTRIP13) indicating the characteristic AAA+ ATPase motifs, the conserved N-terminal domain (NTD) and the extended N-terminal domain exclusive of yeast Pch2 (NTD^EXT^). The positions and sequences of the NESs identified in this work are depicted (purple boxes). **(F)** Fluorescence microscopy images of the localization of GFP-Pch2, GFP-Pch2-nes4A, GFP-NES^TRIP13^-Pch2-nes4A, and GFP-nes7A^TRIP13^-Pch2-nes4A in *zip1Δ* cells. Images were taken at 16 h after meiotic induction. Representative cells are shown. A schematic representation of the different GFP-Pch2 versions is also shown on top. Scale bar, 2 μm. The strain is DP1405 (*zip1Δ pch2Δ*) transformed with the centromeric plasmids pSS393 (*GFP-PCH2*), pSS459 (*GFP-pch2-nes4A*), pSS472 (*GFP-NES^TRIP13^-pch2-nes4A*) and pSS474 (*GFP-nes7A^TRIP13^-pch2-nes4A*). **(G)** Quantification of the ratio of nuclear (including nucleolar) to cytoplasmic GFP fluorescent signal for the experiment shown in (F). Error bars, SD. **(H)** Time course analysis of meiotic nuclear divisions. The percentage of cells containing two or more nuclei is represented. Error bars: SD; n=3. At least 300 cells were scored for each strain at every time point. Strains are: DP1625 (*zip1Δ GFP-PCH2*), DP1986 (*zip1Δ GFP-pch2-nes4*), DP2025 (*zip1Δ GFP-NES^TRIP13^-pch2-nes4A*) and DP2033 (*zip1Δ GFP-nes7A^TRIP13^-pch2-nes4A*). **(I)** Western blot analysis of Hop1-T318 phosphorylation in the strains analyzed in (H) at the indicated time points in meiosis. Pgk1 was used as a loading control. The graph shows the quantification of Hop1-T318ph relative levels.

### Evolutionary conservation of Pch2/TRIP13 nuclear export mechanism

Although the budding yeast Pch2 is meiosis specific, the mammalian homolog TRIP13 is also expressed in somatic cells besides the germline. TRIP13 has been reported to be localized both in the nucleus and the cytoplasm in mammalian somatic cells (Tipton *et al*, 2012; Wang *et al*, 2014; Thul *et al*, 2017). The Pch2/TRIP13 protein family possesses a highly conserved C-terminal AAA+ ATPase domain and a non-catalytic NTD with a lower similarity degree. In addition, yeast Pch2 contains a non-conserved extension at the very beginning of the protein (Figure 4E, Figure S4) that may be involved in the exclusive Pch2 nucleolar localization found in budding yeast via Orc1 interaction (Herruzo *et al*., 2019; Villar-Fernández *et al*., 2020). Using the LocNES prediction tool to analyze the human TRIP13 NTD, we detected a presumptive NES (at positions 65-80) close to the region corresponding to NES^Pch2^ (Figure 4E, S4). To determine whether this putative NES^TRIP13^ is functionally active, we fused it to the NES^Pch2^-deficient *pch2-nes4A* mutant generating a *GFP-NES^TRIP13^-pch2-nes4A* construct. Remarkably, addition of the proposed NES^TRIP13^ partially restored cytoplasmic localization of Pch2-nes4 (Figure 4F, 4G). Likewise, the *zip1Δ GFP-NES^TRIP13^-pch2-nes4A* double mutant displayed a notable meiotic block (Figure 4H) and increased levels of Hop1-T318ph at the 48-h time point (Figure 4I), consistent with a restoration of checkpoint activity. Furthermore, mutation to alanine of all NES^TRIP13^ hydrophobic residues (*nes7A^TRIP13^*) fail to reinstate cytoplasmic localization and checkpoint activity in *pch2-nes4A* (Figure 4F-4I). In sum, these observations strongly suggest that, indeed, the 65-80 amino acid fragment of TRIP13 NTD (**FL**TRN**V**QS**V**S**II**DTE**L**) contains a functional NES capable of substituting the requirement for NES^Pch2^ to implement a balanced localization of Pch2 in the different subcellular compartments during yeast meiosis.

### Concluding remarks

Homeostatic control of Pch2 subcellular localization is crucial for a proper meiotic recombination checkpoint response (Raina & Vader, 2020; Herruzo *et al*., 2021). We unveil here another layer of control of Pch2 function during meiosis involving the nuclear export via the exportin pathway that is essential to maintain the critical balance of Pch2 distribution among different compartments (Figure 5). Functions for Pch2 outside the nucleus are not exclusive of yeast meiosis; in plants, PCH2 also operates in the cytoplasm promoting the nuclear targeting of the HORMA protein ASY1 (Hop1 homolog), in addition to its nuclear action in meiotic chromosome axis remodeling (Balboni *et al*., 2020; Yang *et al*., 2020). We provide evidence here for a possible evolutionary conservation of the nuclear export mechanism for Pch2/TRIP13 from yeast to mammals. In mice, TRIP13 is required for proper completion of meiotic chromosome synapsis and recombination, and *Trip13*-deficient mice are sterile (Li & Schimenti, 2007; Roig *et al*, 2010). Although the localization of TRIP13 during mammalian meiosis has not been reported, it can be detected both in the cytoplasm and nucleus of mouse spermatocytes (I. Roig, personal communication), suggesting that the meiotic role of mammalian TRIP13 may also be modulated by nucleocytoplasmic transport. In humans, biallelic mutations in the *Trip13* gene are associated with female infertility and Wilms tumor (Yost *et al*, 2017; Zhang *et al*, 2020; Hu *et al*, 2022). Interestingly, the CRM1/XPO1 exportin has been recently identified in a screen for potential druggable targets involving TRIP13 function in Wilms tumor derived lines. Furthermore, the FDA-approved drug selinexor (KPT-330), which inhibits nuclear export, leads to suppression of TRIP13 function in these cell lines and it has been proposed as a potential therapeutic strategy for Wilms tumor patients (Mittal *et al*, 2022). In sum, all these observations underscore the importance of Pch2/TRIP13 nucleocytoplasmic traffic in the multiple biological processes impacted by this protein family.

**Figure 5.**
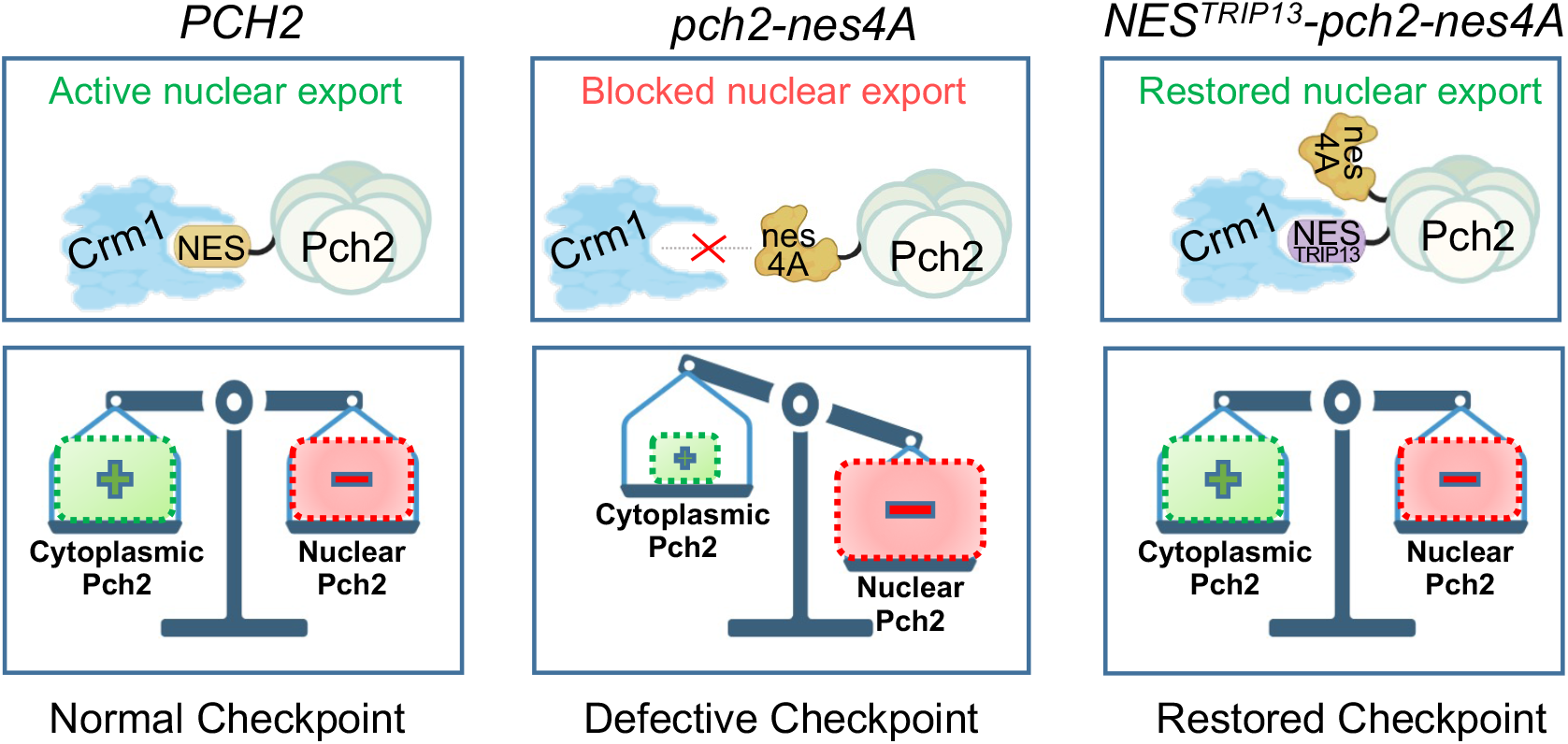
NES-mediated nuclear export of Pch2 is essential for meiotic recombination checkpoint function. The nucleocytoplasmic transport of Pch2 via the exportin pathway maintains Pch2 homeostasis and supports proper checkpoint activity. In the NES-deficient *pch2-nes4A* mutant, the accumulation of Pch2 in the nucleus leads to checkpoint inactivation. The addition of the putative NES of the human TRIP13 ortholog to Pch2-nes4A restores the balanced subcellular distribution of Pch2 and restores checkpoint function. For simplicity, the NES is drawn only in one subunit of the Pch2 hexamer. BioRender.com was used to create this figure.

## MATERIALS AND METHODS

### Yeast strains

The genotypes of yeast strains are listed in Table S1. All strains are in the BR1919 background (Rockmill and Roeder, 1990). The *zip1Δ::LEU2, zip1Δ::LYS2*, *ndt80Δ::kanMX6, pch2Δ::TRP1* and *pch2Δ::URA3* gene deletions were previously described (Sym *et al*., 1993; Sym & Roeder, 1994; San-Segundo & Roeder, 1999; Herruzo *et al*., 2016). The *orc1-3mAID* and *P_HOP1_-GFP-PCH2* constructs have been previously described (Herruzo *et al*., 2019; Herruzo *et al*., 2021). For the experiments involving the inactivation of the Crm1 exportin with LMB we employed diploid strains with both copies of the *CRM1* essential gene deleted (*crm1Δ::hphMX4* or *crm1::natMX4*) using a PCR-based approach (Goldstein & McCusker, 1999) carrying a centromeric plasmid expressing the *crm1-T539C* allele (pSS416) as the only source of exportin in the cell. The *P_HOP1_-GFP-pch2-nes4A, P_HOP1_-GFP-NES^PKI^-pch2-nes4A, P_HOP1_-GFP-NES^TRIP13^-pch2-nes4A,* and *P_HOP1_-GFP-nes7A^TRIP13^-pch2-nes4A* constructs were introduced into the genomic locus of *PCH2* using an adaptation of the *delitto perfetto* technique (Stuckey *et al*, 2011). Basically, PCR fragments flanked by the appropriate sequences containing the *HOP1* promoter followed by the *GFP-pch2-nes4A*, *GFP-NES^PKI^-pch2-nes4A, GFP-NES^TRIP13^-pch2-nes4A* or *GFP-nes7A^TRIP13^-pch2-nes4A* sequences, and a five Gly-Ala repeat linker before the second codon of *PCH2,* were amplified from pSS459, pSS462, pSS472 and pSS474, respectively (see below). These fragments were transformed into a strain carrying the CORE cassette (*kanMX4-URA3*) inserted close to the 5’ end of *PCH2.* G418-sensitive and 5-FOA resistant clones containing the correct integrated construct, which results in the elimination of 91 nt of the *PCH2* promoter, were selected. All constructions and mutations were verified by PCR analysis and/or sequencing. The sequences of all primers used in strain construction are available upon request. All strains were made by direct transformation of haploid parents or by genetic crosses always in an isogenic background. Diploids were made by mating the corresponding haploid parents and isolation of zygotes by micromanipulation.

### Plasmids

The plasmids used are listed in Table S2. The pSS393 centromeric plasmid expressing *P_HOP1_-GFP-PCH2* was previously described (Herruzo *et al*., 2019). The pSS416 plasmid containing *crm1-T539C* was kindly provided by M. Dosil (CIC, Salamanca). The pSS448 and pSS451 plasmids, driving the expression of *P_HOP1_-GFP-pch2^(98-107)-6A^* and *P_HOP1_-GFP-pch2^(127-136)-5A^,* respectively, were made by non-overlapping mutagenesis following the procedure described in the Q5 site-directed mutagenesis kit (New England Biolabs), using the pSS393 as template and divergent primers. The forward primers carried the sequence encoding the mutated 98-107 (**LI**RS**L**AK**VLL** to **AA**RS**A**AK**AAA**) or 127-136 (**L**F**L**S**L**F**V**KK**I** to **A**F**A**S**A**F**A**KK**A**) regions in the Pch2 NTD. The pSS459 plasmid driving the expression of *P_HOP1-_GFP-pch2-nes4A* was derived from pSS393 by using the NEBuilder assembly kit (New England Biolabs) and a synthesized gBlock fragment (IDT) containing the mutated sequence corresponding to the 205-214 region (**L**STE**F**DK**I**D**L** to **A**STE**A**DK**A**D**A**). To construct the pSS462 plasmid containing *P_HOP1-_GFP-NES^PKI^-pch2-nes4A*, a 1.8-kb fragment was amplified from pSS459 (*P_HOP1-_GFP-pch2-nes4A)*, using a forward primer containing the sequence encoding the Nuclear Export Signal (NES) from the PKI protein (LALKLAGLDI) (Wen *et al*., 1995) preceded by a *Not*I site at the 5’ end and a reverse primer within the *PCH2* coding sequence downstream of the endogenous *Blp*I site. The fragment was digested with *Not*I-*Blp*I and cloned into the same sites of pSS393. To generate the pSS472 plasmid that contains *P_HOP1-_GFP-NES^TRIP13^-pch2-nes4A*, a 418-bp fragment was amplified from pSS393 (*P_HOP1-_GFP-PCH2*) using a forward primer containing the presumptive NES of TRIP13 (at positions 65-80) preceded by a *Not*I site at its 5’end, and a reverse primer within the *PCH2* coding sequence downstream of the endogenous *BamH*I site. The fragment was digested with *Not*I-*BamH*I and cloned into the same sites of pSS459. To make the pSS474 plasmid containing *P_HOP1-_GFP-nes7A^TRIP13^-pch2-nes4A*, a 418pb fragment was amplified from pSS393, using a forward primer containing the mutation to alanine of all NES^TRIP13^ hydrophobic residues (**FL**TRN**V**QS**V**S**II**DTE**L** to **AA**TRN**A**QS**A**S**AA**DTE**A**), preceded by a *Not*I site at its 5’end, and a reverse primer within the *PCH2* coding sequence downstream of the endogenous *BamH*I site. The fragment was digested with *Not*I-*BamH*I and cloned into the same sites of pSS459.

### Meiotic cultures and meiotic time courses

To induce meiosis and sporulation, BR strains were grown in 3.5 ml of synthetic complete medium (2% glucose, 0.7% yeast nitrogen base without amino acids, 0.05% adenine, and complete supplement mixture from Formedium at twice the particular concentration indicated by the manufacturer) for 20–24 h, then transferred to 2.5 ml of YPDA (1% yeast extract, 2% peptone, 2% glucose, and 0.02% adenine) and incubated to saturation for an additional 8 h. Cells were harvested, washed with 2% potassium acetate (KAc), resuspended into 2% KAc (10 ml), and incubated at 30°C with vigorous shaking to induce meiosis. Both YPDA and 2% KAc were supplemented with 20 mM adenine and 10 mM uracil. The culture volumes were scaled up when needed. To inhibit Crm1-T539C, cultures were treated with 500 ng/ml of leptomycin B (LMB) at 15h in meiosis for the indicated periods of time. To induce Orc1-3mAID degradation, auxin (500μM) was added to the cultures 12 h after meiotic induction. To score meiotic nuclear divisions, samples from meiotic cultures were taken at different time points, fixed in 70% ethanol, washed in phosphate-buffered saline (PBS) and stained with 1 μg/μl 4′,6-diamidino-2-phenylindole (DAPI) for 15 min. At least 300 cells were counted at each time point. Meiotic time courses were repeated several times; averages and error bars from at least three replicates are shown.

### Western blotting

Total cell extracts for Western blot analysis were prepared by trichloroacetic acid (TCA) precipitation from 3-ml aliquots of sporulation cultures, as previously described (Acosta *et al*, 2011). The antibodies used are listed in Table S3. The Pierce ECL Plus reagents (ThermoFisher Scientific) were used for detection. The signal was captured with a Fusion FX6 system (Vilber) and quantified with the Evolution-Capt software (Vilber).

### Cytology

Immunofluorescence of chromosome spreads was performed essentially as described (Rockmill, 2009). The antibodies used are listed in Table S3. Images of spreads were captured with a Nikon Eclipse 90i fluorescence microscope controlled with MetaMorph software (Molecular Devices) and equipped with a Hammamatsu Orca-AG charge-coupled device (CCD) camera and a PlanApo VC 100x 1.4 NA objective. To measure Pch2 and Hop1 intensity on chromosome spreads, a region containing DAPI-stained chromatin was defined and the Raw Integrated Density values were measured. Background values were subtracted using the rolling ball algorithm from Fiji setting the radius to 50 pixels. Images of whole live cells expressing *GFP-PCH2* were captured with an Olympus IX71 fluorescence microscope equipped with a personal DeltaVision system, a CoolSnap HQ2 (Photometrics) camera, and 100x UPLSAPO 1.4 NA objective. Stacks of 7 planes at 0.8-μm intervals were collected. Maximum intensity projections of 3 planes containing GFP-Pch2 are shown for *ZIP1* cells and single planes for *zip1Δ* cells. To determine the nuclear/cytoplasm GFP fluorescence ratio shown in Figures 1D, 3B, 4B, 4G and S3B, the ROI manager tool of Fiji software (Schindelin *et al*, 2012) was used to define the cytoplasm and nuclear (including the nucleolus) areas; the mean intensity values were measured and subjected to background subtraction. In the specific case of Figure 4G, due to the difficulty in the discrimination of the nuclei in the strain expressing *GFP-NES^TRIP13^-pch2-nes4A*, we measured the average area of more than 50 nuclei from each one of the other strains analyzed in that Figure in which the contour of the nucleus was visible, resulting in a value of 7.02 μm^2^. We then applied this value to draw a circle containing the nucleolus in all the cells to define the nucleus and measured the nuclear/cytoplasmic fluorescence intensity ratio.

### Sporulation efficiency

Sporulation efficiency was quantitated by microscopic examination of asci formation after 3 days on sporulation plates. Both mature and immature asci were scored. At least 300 cells were counted for every strain.

### Statistics

To determine the statistical significance of differences, a two-tailed Student t-test was used. *P*-values were calculated with the GraphPad Prism 9.0 software. *P*<0.05 (*); *P*<0.01 (**); *P*<0.001 (***); *P*<0.0001 (****). The nature of the error bars in the graphical representations and the number of biological replicates are indicated in the corresponding figure legend.

## ACKNOWLEDGEMENTS

We are grateful to Andrés Clemente for helpful comments and discussions and to Ignasi Roig for sharing unpublished results. We also thank Mercedes Dosil and Yolanda Sánchez for reagents, Jesús Pinto, Carmen Castro and Carlos R. Vázquez for advice on microscopy analysis, Isabel Acosta for technical support, and Ana Lago-Maciel for help in strains construction. This work was supported by the grant PID2021-125830NB-I00 from Ministry of Science and Innovation (MCIN/AEI/FEDER, EU) of Spain to PSS and JAC. EH was partially supported by the grant CSI259P20 from the “Junta de Castilla y León” (FEDER, EU). ESD is supported by a predoctoral contract from the “Junta de Castilla y León” (co-funded by the Education Department and the European Social Fund FSE+).

## AUTHOR CONTRIBUTIONS

EH: conceptualization, investigation, formal analysis, visualization. ESD: conceptualization, investigation, formal analysis, visualization. SGA: investigation, formal analysis.

BS: conceptualization, supervision, project administration. JAC: conceptualization, resources, funding acquisition

PSS: conceptualization, investigation, supervision, funding acquisition, visualization, writing original draft.

All authors revised, commented, and approved the manuscript.

## Supplemental Figure Legends

**Figure S1.**
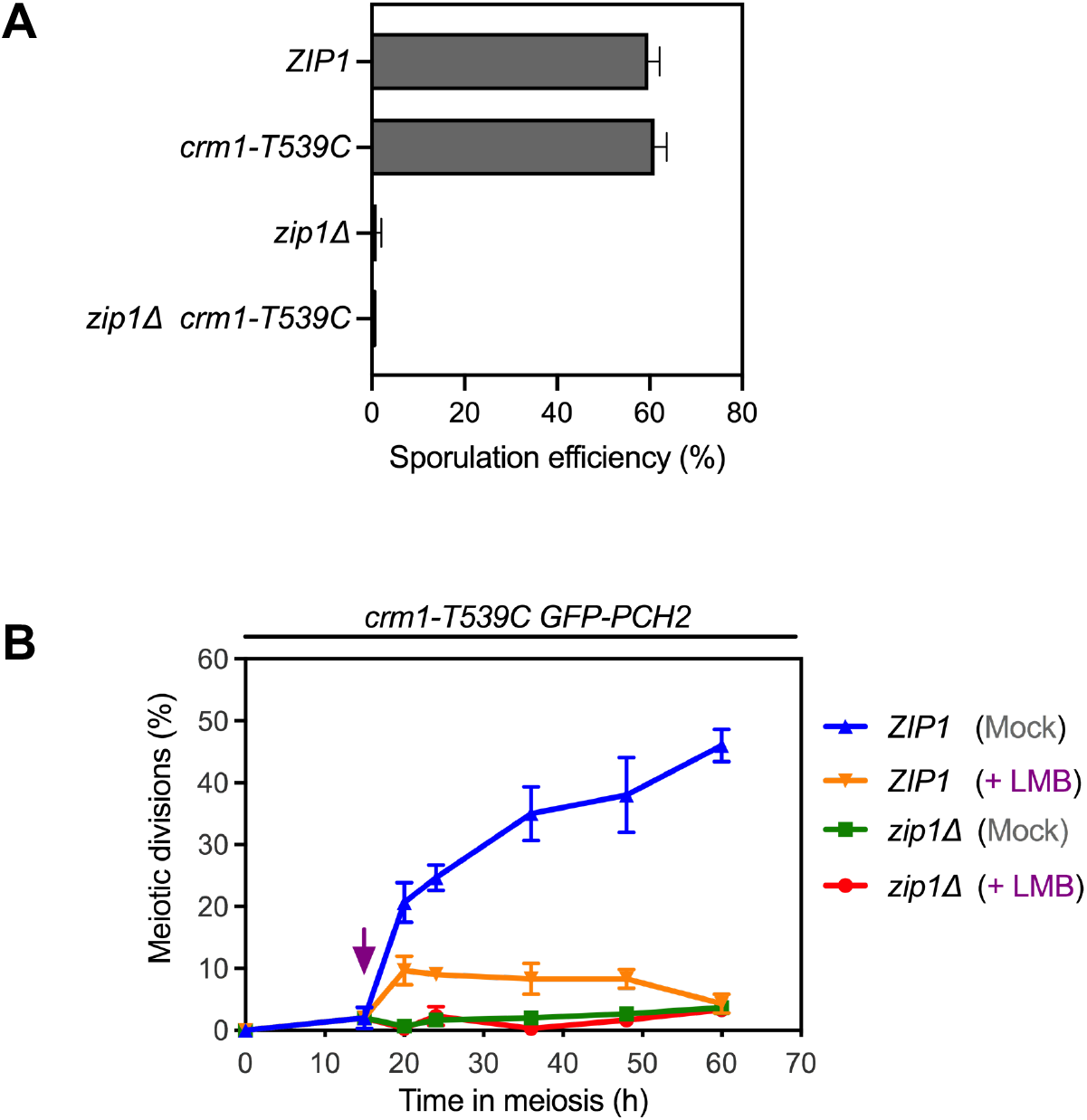
The *crm1-T539C* mutation does not substantially impair meiosis or checkpoint function in the absence of LMB. **(A)** Sporulation efficiency was examined after 3 days on sporulation plates. Error bars, SD; n=3. At least 300 cells were counted for each strain. Strains are: DP421 (wild type), DP1717 (*crm1-T539C GFP-PCH2*), DP422 (*zip1Δ*), and DP1721 (*zip1Δ crm1-T539C GFP-PCH2*). **(B)** Time course analysis of meiotic nuclear divisions. The percentage of cells containing two or more nuclei is represented. Ethanol (Mock) or Leptomycin B (LMB) were added 15 h after meiotic induction (arrow). Error bars: SD; n=3. At least 300 cells were scored for each strain at every time point. Strains are: DP1717 (*crm1-T539C GFP-PCH2*) and DP1721 (*zip1Δ crm1-T539C GFP-PCH2*).

**Figure S2.**
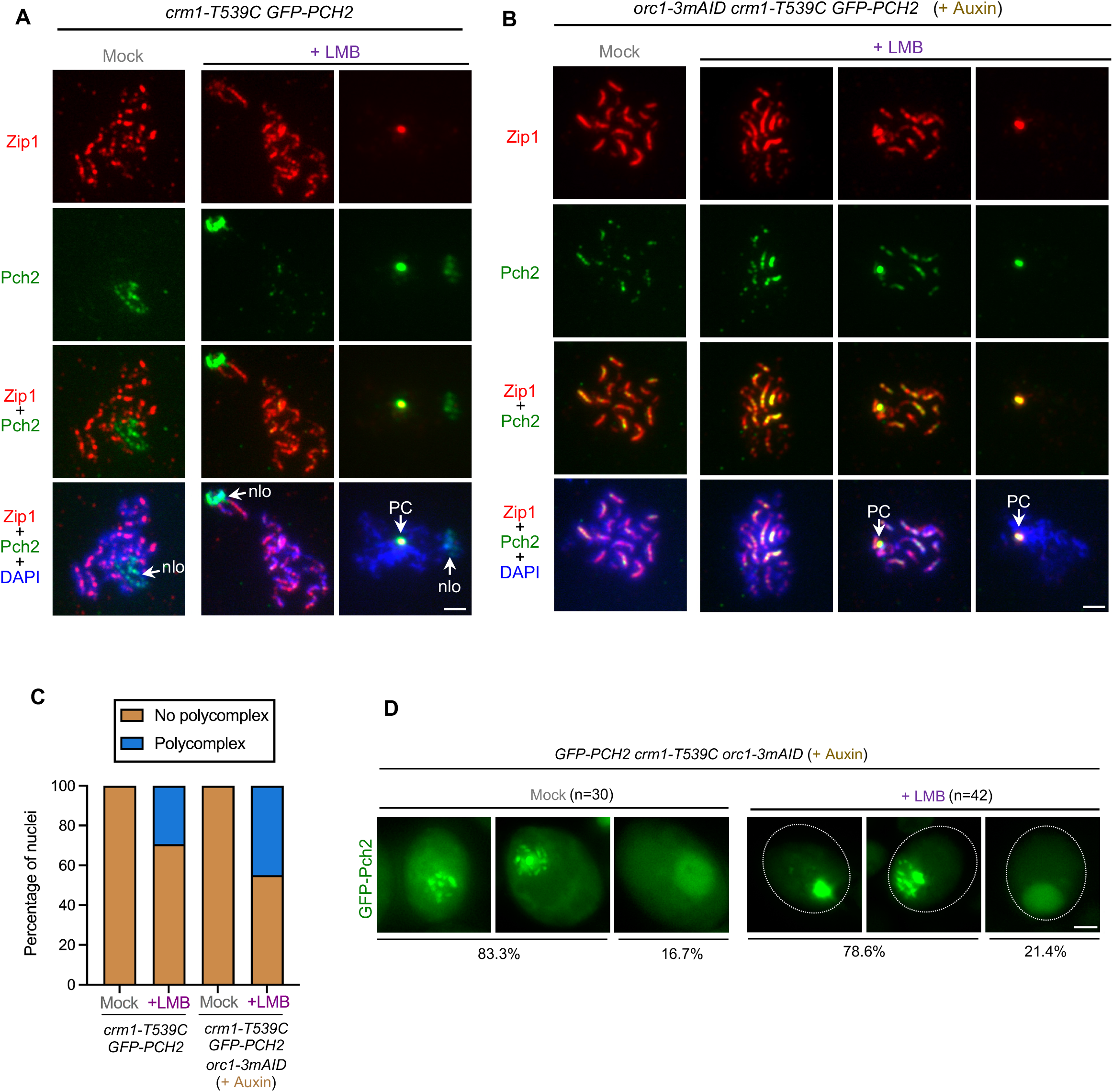
Nuclear accumulation of Pch2 is linked to increased association of Pch2 with the SC and SC assemblies in *ZIP1+* cells. **(A-B)** Immunofluorescence of spread meiotic chromosomes at pachytene stained with anti-GFP antibodies (to detect GFP-Pch2; green), anti-Zip1 antibodies (red) and DAPI (blue). Representative nuclei are shown. In both, (A) and (B), cultures were mock-treated, or treated with 500 ng/ml LMB 15 h after meiotic induction. In (B), Auxin (500μM) was also added 12 h after meiotic induction to degrade Orc1. Spreads were prepared at 19 h. Arrows point to the rDNA region (nlo) and Polycomplex (PC). Scale bar, 2 μm. The strain in (A) is: DP1717 (*crm1-T539C GFP-PCH2*). The strain in (B) is: DP1885 (*orc1-3mAID crm1-T539C GFP-PCH2*). **(C)** Percentage of nuclei containing polycomplexes in the experiments shown in A and B. Between 10 to 20 nuclei were counted for each strain and condition. **(D)** Quantification of the different patterns of Pch2 localization in the experiment presented in Figure 3A-3B. Representative cells are shown.

**Figure S3.**
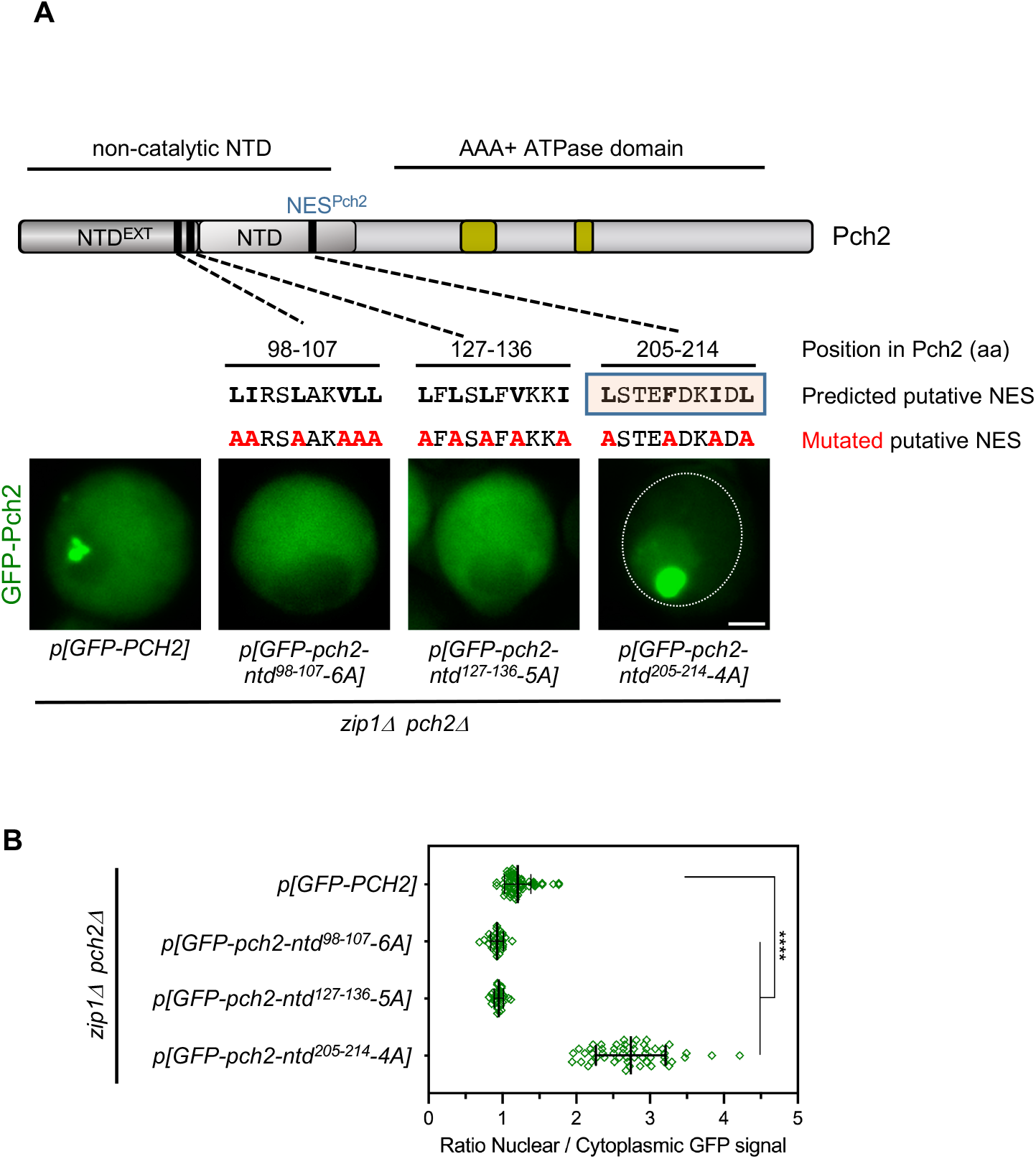
Identification of NES sequences in Pch2. **(A)** Schematic representation of the *S. cerevisiae* Pch2 protein. The position and sequence of the three putative NESs predicted by LocNES in the non-catalytic N-terminal domain of Pch2 are depicted, as well as the corresponding mutants generated. The images show representative *zip1Δ pch2Δ* cells transformed with centromeric plasmids expressing wild-type *GFP-PCH2* or the different mutated versions of the predicted NESs, as indicated. Note that only the mutation of the 205-214 region (boxed) leads to Pch2 accumulation in the nucleus. Images were taken 15 h after meiotic induction. The strain is DP1405 (*zip1Δ pch2Δ*) transformed with the centromeric plasmids pSS393 (*GFP-PCH2*), pSS448 (*GFP-pch2-ntd^98-107^-6A*), pSS451 (*GFP-pch2-ntd^127-136^-5A*) and pSS459 (*GFP-pch2-ntd^205-214^-4A*). **(B)** Quantification of the ratio of nuclear (including nucleolar) to cytoplasmic GFP fluorescent signal for the experiment shown in (A). Error bars, SD.

**Figure S4.**
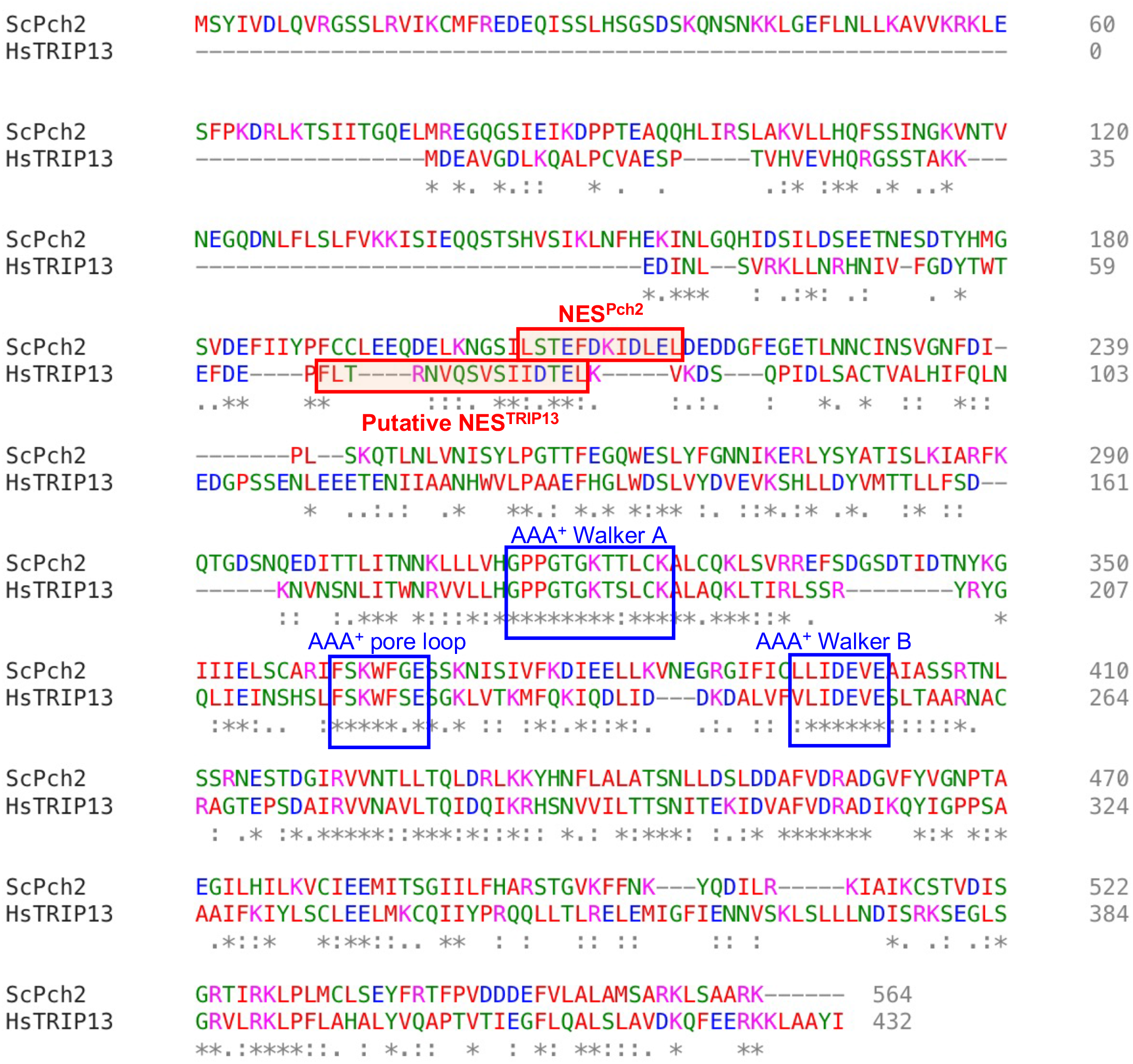
Conserved position of the NES in Pch2 and TRIP13. ClustalW alignment of the protein sequences of Pch2 orthologs from *S. cerevisiae* (ScPch2) and human (HsTRIP13). The characteristic AAA+ ATPase features are boxed in blue. The identified NESs are boxed in red. The color code for amino acids is the following: AVFPMILW (small + hydrophobic -Y): red. DE (acidic): blue. RK (basic -H): magenta. STYHCNGQ (hydroxyl +sulfhydryl + amine + G): green. Alignment was performed at https://www.ebi.ac.uk/Tools/msa/clustalo/.

---

**Table S1.**
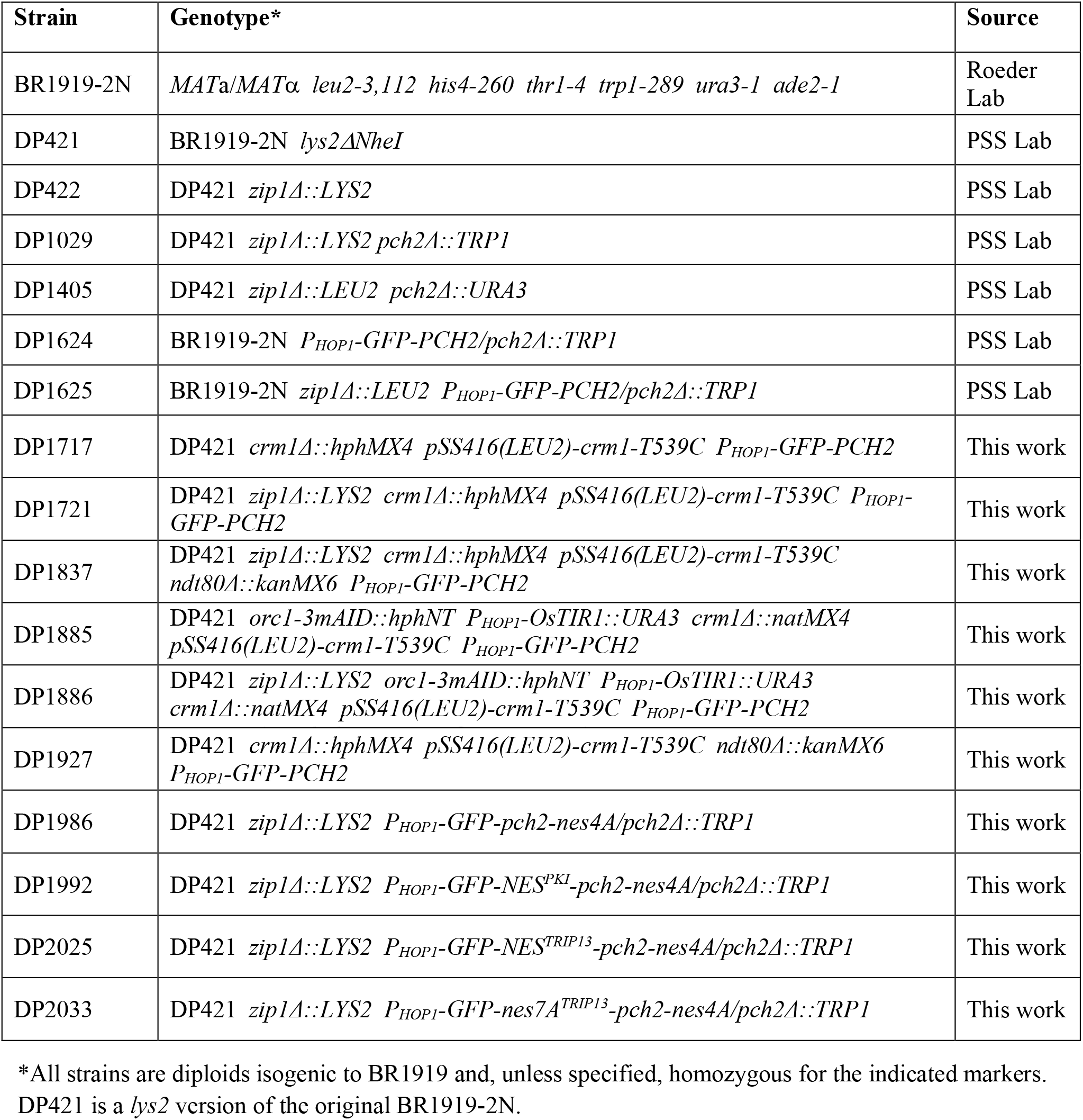
Saccharomyces cerevisiae strains.

**Table S2.**
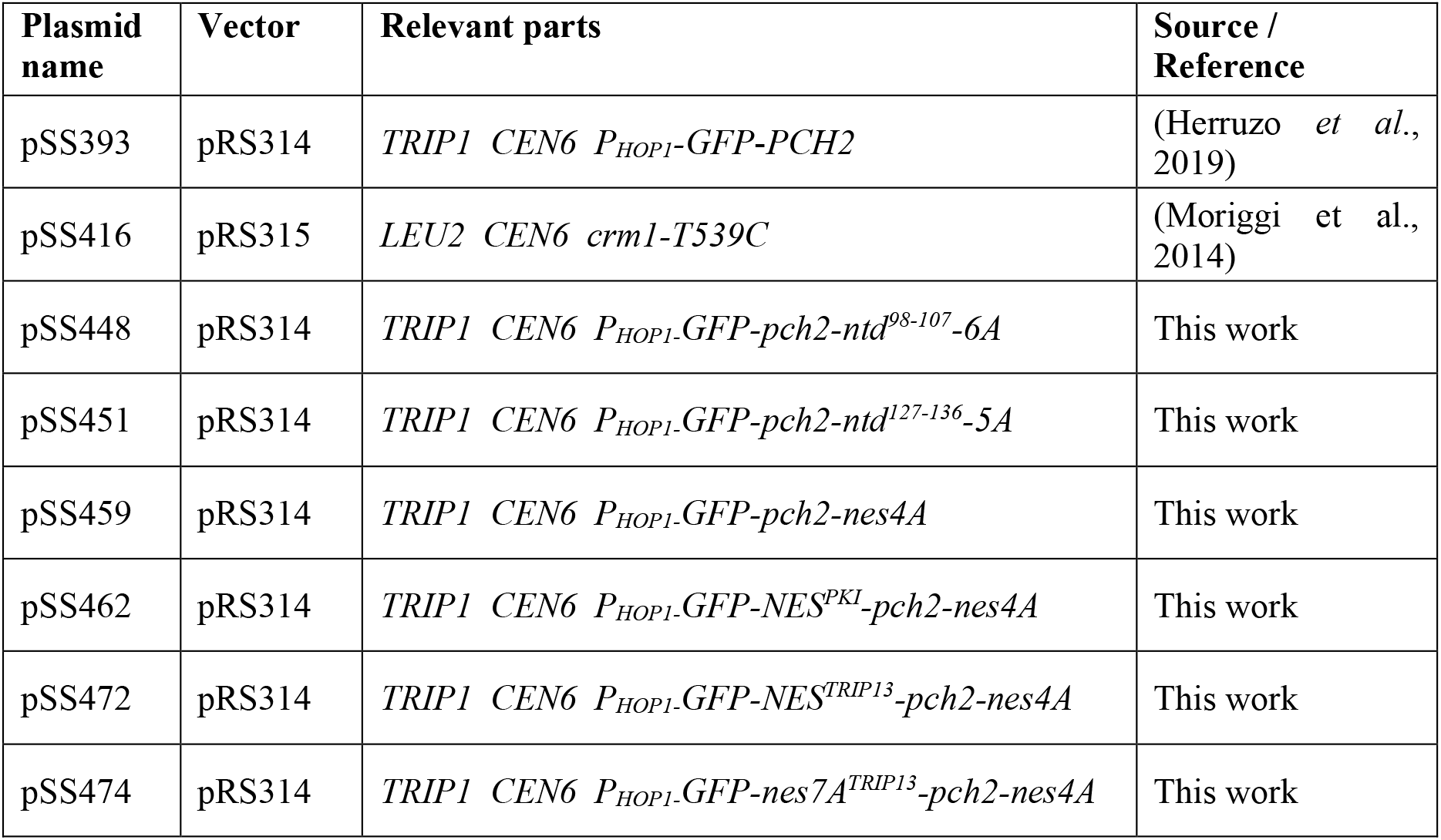
Plasmids.

**Table S3.** Primary antibodies.

| Antibody | Host and type | Application*<br>(Dilution) | Source / Reference |
| --- | --- | --- | --- |
| Hop1 | Rabbit polyclonal | IF (1:300) | (1) |
| Hop1-T318-P | Rabbit polyclonal | WB (1:1000) | (2) |
| Pgk1 (22C5D8) | Mouse monoclonal | WB (1:5000) | Molecular Probes<br>459250 |
| Pch2 | Rabbit polyclonal | WB (1:2000)<br>IF (1:200) | (3) |
| Nsr1 (31C4) | Mouse monoclonal | IF (1:200) | ThermoFisher<br>MA1-10030 |
| Zip1 | Rabbit polyclonal | IF (1:200) | (4) |
| GFP (JL-8) | Mouse monoclonal | IF (1:200) | Clontech<br>632381 |
| Cdc5 | Goat polyclonal | WB (1:1000) | Santa Cruz Biotechnology<br>sc-6733 |

\*WB, western blot; IF, immunofluorescence

## REFERENCES

Acosta I, Ontoso D, San-Segundo PA (2011) The budding yeast polo-like kinase Cdc5 regulates the Ndt80 branch of the meiotic recombination checkpoint pathway. Mol Biol Cell 22: 3478–3490

Alfieri C, Chang L, Barford D (2018) Mechanism for remodelling of the cell cycle checkpoint protein MAD2 by the ATPase TRIP13. Nature 559: 274–278

Balboni M, Yang C, Komaki S, Brun J, Schnittger A (2020) COMET functions as a PCH2 cofactor in regulating the HORMA domain protein ASY1. Curr Biol 30: 4113–4127.e4116

Bhalla N (2023) PCH-2 and meiotic HORMADs: A module for evolutionary innovation in meiosis? Curr Top Dev Biol 151: 317–344

Bhalla N, Dernburg AF (2005) A conserved checkpoint monitors meiotic chromosome synapsis in *Caenorhabditis elegans*. Science 310: 1683–1686

Bolcun-Filas E, Handel MA (2018) Meiosis: the chromosomal foundation of reproduction. Biol Reprod 99: 112–126

Borner GV, Barot A, Kleckner N (2008) Yeast Pch2 promotes domainal axis organization, timely recombination progression, and arrest of defective recombinosomes during meiosis. Proc Natl Acad Sci USA 105: 3327–3332

Callender TL, Laureau R, Wan L, Chen X, Sandhu R, Laljee S, Zhou S, Suhandynata RT, Prugar E, Gaines WA et al (2016) Mek1 down regulates Rad51 activity during yeast meiosis by phosphorylation of Hed1. PLoS Genet 12: e1006226

Carballo JA, Johnson AL, Sedgwick SG, Cha RS (2008) Phosphorylation of the axial element protein Hop1 by Mec1/Tel1 ensures meiotic interhomolog recombination. Cell 132: 758–770

Cardoso da Silva R, Villar-Fernández MA, Vader G (2020) Active transcription and Orc1 drive chromatin association of the AAA+ ATPase Pch2 during meiotic G2/prophase. PLoS Genet 16: e1008905

Cavero S, Herruzo E, Ontoso D, San-Segundo PA (2016) Impact of histone H4K16 acetylation on the meiotic recombination checkpoint in *Saccharomyces cerevisiae*. Microb Cell 3: 606–620

Chen C, Jomaa A, Ortega J, Alani EE (2014) Pch2 is a hexameric ring ATPase that remodels the chromosome axis protein Hop1. Proc Natl Acad Sci USA 111: E44–53

Chen X, Gaglione R, Leong T, Bednor L, de Los Santos T, Luk E, Airola M, Hollingsworth NM (2018) Mek1 coordinates meiotic progression with DNA break repair by directly phosphorylating and inhibiting the yeast pachytene exit regulator Ndt80. PLoS Genet 14: e1007832

Dong H, Roeder GS (2000) Organization of the yeast Zip1 protein within the central region of the synaptonemal complex. J Cell Biol 148: 417–426

Eytan E, Wang K, Miniowitz-Shemtov S, Sitry-Shevah D, Kaisari S, Yen TJ, Liu ST, Hershko A (2014) Disassembly of mitotic checkpoint complexes by the joint action of the AAA-ATPase TRIP13 and p31(comet). Proc Natl Acad Sci U S A 111: 12019–12024

Farmer S, Hong EJ, Leung WK, Argunhan B, Terentyev Y, Humphryes N, Toyoizumi H, Tsubouchi H (2012) Budding yeast Pch2, a widely conserved meiotic protein, is involved in the initiation of meiotic recombination. PLoS One 7: e39724

Fung HY, Fu SC, Chook YM (2017) Nuclear export receptor CRM1 recognizes diverse conformations in nuclear export signals. Elife 6: e23961

Fung HYJ, Niesman A, Chook YM (2021) An update to the CRM1 cargo/NES database NESdb. Mol Biol Cell 32: 467–469

Giacopazzi S, Vong D, Devigne A, Bhalla N (2020) PCH-2 collaborates with CMT-1 to proofread meiotic homolog interactions. PLoS Genet 16: e1008904

Goldstein AL, McCusker JH (1999) Three new dominant drug resistance cassettes for gene disruption in *Saccharomyces cerevisiae*. Yeast 15: 1541–1553

González-Arranz S, Gardner JM, Yu Z, Patel NJ, Heldrich J, Santos B, Carballo JA, Jaspersen SL, Hochwagen A, San-Segundo PA (2020) SWR1-independent association of H2A.Z to the LINC complex promotes meiotic chromosome motion. Front Cell Dev Biol 8: 594092

Hanson PI, Whiteheart SW (2005) AAA+ proteins: have engine, will work. Nat Rev Mol Cell Biol 6: 519–529

Heldrich J, Sun X, Vale-Silva LA, Markowitz TE, Hochwagen A (2020) Topoisomerases modulate the timing of meiotic DNA breakage and chromosome morphogenesis in *Saccharomyces cerevisiae*. Genetics 215: 59–73

Herruzo E, Lago-Maciel A, Baztan S, Santos B, Carballo JA, San-Segundo PA (2021) Pch2 orchestrates the meiotic recombination checkpoint from the cytoplasm. PLoS Genet 17: e1009560

Herruzo E, Ontoso D, González-Arranz S, Cavero S, Lechuga A, San-Segundo PA (2016) The Pch2 AAA+ ATPase promotes phosphorylation of the Hop1 meiotic checkpoint adaptor in response to synaptonemal complex defects. Nucleic Acids Res 44: 7722–7741

Herruzo E, Santos B, Freire R, Carballo JA, San-Segundo PA (2019) Characterization of Pch2 localization determinants reveals a nucleolar-independent role in the meiotic recombination checkpoint. Chromosoma 128: 297–316

Hollingsworth NM, Gaglione R (2019) The meiotic-specific Mek1 kinase in budding yeast regulates interhomolog recombination and coordinates meiotic progression with double-strand break repair. Curr Genet 65: 631–641

Hollingsworth NM, Goetsch L, Byers B (1990) The *HOP1* gene encodes a meiosis-specific component of yeast chromosomes. Cell 61: 73–84

Hu H, Zhang S, Guo J, Meng F, Chen X, Gong F, Lu G, Zheng W, Lin G (2022) Identification of Novel Variants of Thyroid Hormone Receptor Interaction Protein 13 That Cause Female Infertility Characterized by Zygotic Cleavage Failure. Front Physiol 13: 899149

Humphryes N, Leung WK, Argunhan B, Terentyev Y, Dvorackova M, Tsubouchi H (2013) The Ecm11-Gmc2 complex promotes synaptonemal complex formation through assembly of transverse filaments in budding yeast. PLoS Genet 9: e1003194

Hunter N (2015) Meiotic Recombination: The Essence of Heredity. Cold Spring Harb Perspect Biol 7: a016618

Joshi N, Barot A, Jamison C, Borner GV (2009) Pch2 links chromosome axis remodeling at future crossover sites and crossover distribution during yeast meiosis. PLoS Genet 5: e1000557

Joshi N, Brown MS, Bishop DK, Borner GV (2015) Gradual implementation of the meiotic recombination program via checkpoint pathways controlled by global DSB levels. Mol Cell 57: 797–811

Joyce EF, McKim KS (2009) Drosophila PCH2 is required for a pachytene checkpoint that monitors double-strand-break-independent events leading to meiotic crossover formation. Genetics 181: 39–51

Keeney S, Lange J, Mohibullah N (2014) Self-organization of meiotic recombination initiation: general principles and molecular pathways. Annu Rev Genet 48: 187–214

Kim HJ, Liu C, Dernburg AF (2022) How and Why Chromosomes Interact with the Cytoskeleton during Meiosis. Genes 13: 901

Klein F, Mahr P, Galova M, Buonomo SB, Michaelis C, Nairz K, Nasmyth K (1999) A central role for cohesins in sister chromatid cohesion, formation of axial elements, and recombination during yeast meiosis. Cell 98: 91–103

Lambing C, Osman K, Nuntasoontorn K, West A, Higgins JD, Copenhaver GP, Yang J, Armstrong SJ, Mechtler K, Roitinger E et al (2015) Arabidopsis PCH2 Mediates Meiotic Chromosome Remodeling and Maturation of Crossovers. PLoS Genet 11: e1005372

Láscarez-Lagunas L, Martinez-Garcia M, Colaiacovo M (2020) SnapShot: Meiosis – Prophase I. Cell 181: 1442

Li XC, Schimenti JC (2007) Mouse pachytene checkpoint 2 (trip13) is required for completing meiotic recombination but not synapsis. PLoS Genet 3: e130

Lo YH, Chuang CN, Wang TF (2014) Pch2 prevents Mec1/Tel1-mediated Hop1 phosphorylation occurring independently of Red1 in budding yeast meiosis. PLoS One 9: e85687

Mittal K, Lee BP, Cooper GW, Shim J, Jonus HC, Kim WJ, Doshi M, Almanza D, Kynnap BD, Christie AL et al (2022) Targeting TRIP13 in Wilms Tumor with Nuclear Export Inhibitors. bioRxiv: 2022.2002.2023.481521

Neville M, Rosbash M (1999) The NES-Crm1p export pathway is not a major mRNA export route in *Saccharomyces cerevisiae*. EMBO J 18: 3746–3756

Niu H, Wan L, Baumgartner B, Schaefer D, Loidl J, Hollingsworth NM (2005) Partner choice during meiosis is regulated by Hop1-promoted dimerization of Mek1. Mol Biol Cell 16: 5804–5818

Niu H, Wan L, Busygina V, Kwon Y, Allen JA, Li X, Kunz RC, Kubota K, Wang B, Sung P et al (2009) Regulation of meiotic recombination via Mek1-mediated Rad54 phosphorylation. Mol Cell 36: 393–404

Ontoso D, Acosta I, van Leeuwen F, Freire R, San-Segundo PA (2013) Dot1-dependent histone H3K79 methylation promotes activation of the Mek1 meiotic checkpoint effector kinase by regulating the Hop1 adaptor. PLoS Genet 9: e1003262

Penedos A, Johnson AL, Strong E, Goldman AS, Carballo JA, Cha RS (2015) Essential and checkpoint functions of budding yeast ATM and ATR during meiotic prophase are facilitated by differential phosphorylation of a meiotic adaptor protein, Hop1. PLoS One 10: e0134297

Prince JP, Martinez-Perez E (2022) Functions and Regulation of Meiotic HORMA-Domain Proteins. Genes 13: 777

Puchades C, Sandate CR, Lander GC (2020) The molecular principles governing the activity and functional diversity of AAA+ proteins. Nat Rev Mol Cell Biol 21: 43–58

Raina VB, Vader G (2020) Homeostatic control of meiotic prophase checkpoint function by Pch2 and Hop1. Curr Biol 30: 4413–4424 e4415

Refolio E, Cavero S, Marcon E, Freire R, San-Segundo PA (2011) The Ddc2/ATRIP checkpoint protein monitors meiotic recombination intermediates J Cell Sci 124: 2488–2500

Rockmill B (2009) Chromosome spreading and immunofluorescence methods in *Saccharomyes cerevisiae*. Methods Mol Biol 558: 3–13

Roig I, Dowdle JA, Toth A, de Rooij DG, Jasin M, Keeney S (2010) Mouse TRIP13/PCH2 is required for recombination and normal higher-order chromosome structure during meiosis. PLoS Genet 6: e1001062

San-Segundo PA, Clemente-Blanco A (2020) Resolvases, dissolvases, and helicases in homologous recombination: clearing the road for chromosome segregation. Genes 11: 71

San-Segundo PA, Roeder GS (1999) Pch2 links chromatin silencing to meiotic checkpoint control. Cell 97: 313–324

San-Segundo PA, Roeder GS (2000) Role for the silencing protein Dot1 in meiotic checkpoint control. Mol Biol Cell 11: 3601–3615

Schindelin J, Arganda-Carreras I, Frise E, Kaynig V, Longair M, Pietzsch T, Preibisch S, Rueden C, Saalfeld S, Schmid B et al (2012) Fiji: an open-source platform for biological-image analysis. Nat Methods 9: 676–682

Smith AV, Roeder GS (1997) The yeast Red1 protein localizes to the cores of meiotic chromosomes. J Cell Biol 136: 957–967

Stade K, Ford CS, Guthrie C, Weis K (1997) Exportin 1 (Crm1p) is an essential nuclear export factor. Cell 90: 1041–1050

Stuckey S, Mukherjee K, Storici F (2011) In vivo site-specific mutagenesis and gene collage using the delitto perfetto system in yeast *Saccharomyces cerevisiae*. Methods Mol Biol 745: 173–191

Subramanian VV, Hochwagen A (2014) The meiotic checkpoint network: step-by-step through meiotic prophase. Cold Spring Harb Perspect Biol 6: a016675

Subramanian VV, MacQueen AJ, Vader G, Shinohara M, Sanchez A, Borde V, Shinohara A, Hochwagen A (2016) Chromosome synapsis alleviates Mek1-dependent suppression of meiotic DNA repair. PLoS Biol 14: e1002369

Subramanian VV, Zhu X, Markowitz TE, Vale-Silva LA, San-Segundo PA, Hollingsworth NM, Keeney S, Hochwagen A (2019) Persistent DNA-break potential near telomeres increases initiation of meiotic recombination on short chromosomes. Nat Commun 10: 970

Sun Q, Carrasco YP, Hu Y, Guo X, Mirzaei H, Macmillan J, Chook YM (2013) Nuclear export inhibition through covalent conjugation and hydrolysis of Leptomycin B by CRM1. Proc Natl Acad Sci U S A 110: 1303–1308

Sym M, Engebrecht JA, Roeder GS (1993) ZIP1 is a synaptonemal complex protein required for meiotic chromosome synapsis. Cell 72: 365–378

Sym M, Roeder GS (1994) Crossover interference is abolished in the absence of a synaptonemal complex protein. Cell 79: 283–292

Thul PJ, Akesson L, Wiking M, Mahdessian D, Geladaki A, Ait Blal H, Alm T, Asplund A, Bjork L, Breckels LM et al (2017) A subcellular map of the human proteome. Science 356

Tipton AR, Wang K, Oladimeji P, Sufi S, Gu Z, Liu ST (2012) Identification of novel mitosis regulators through data mining with human centromere/kinetochore proteins as group queries. BMC Cell Biol 13: 15

Vader G (2015) Pch2(TRIP13): controlling cell division through regulation of HORMA domains. Chromosoma 124: 333–339

Vader G, Blitzblau HG, Tame MA, Falk JE, Curtin L, Hochwagen A (2011) Protection of repetitive DNA borders from self-induced meiotic instability. Nature 477: 115–119

Villar-Fernández MA, Cardoso da Silva R, Firlej M, Pan D, Weir E, Sarembe A, Raina VB, Bange T, Weir JR, Vader G (2020) Biochemical and functional characterization of a meiosis specific Pch2/ORC AAA+ assembly. Life Sci Alliance 3: e201900630

Wang K, Sturt-Gillespie B, Hittle JC, Macdonald D, Chan GK, Yen TJ, Liu ST (2014) Thyroid hormone receptor interacting protein 13 (TRIP13) AAA-ATPase is a novel mitotic checkpoint silencing protein. J Biol Chem 289: 23928–23937

Wang Y, Chang CY, Wu JF, Tung KS (2011) Nuclear localization of the meiosis-specific transcription factor Ndt80 is regulated by the pachytene checkpoint. Mol Biol Cell 22: 1878–1886

Wen W, Meinkoth JL, Tsien RY, Taylor SS (1995) Identification of a signal for rapid export of proteins from the nucleus. Cell 82: 463–473

West AMV, Komives EA, Corbett KD (2018) Conformational dynamics of the Hop1 HORMA domain reveal a common mechanism with the spindle checkpoint protein Mad2. Nucleic Acids Res 46: 279–292

Xu D, Marquis K, Pei J, Fu SC, Cagatay T, Grishin NV, Chook YM (2015) LocNES: a computational tool for locating classical NESs in CRM1 cargo proteins. Bioinformatics 31: 1357–1365

Yang C, Hu B, Portheine SM, Chuenban P, Schnittger A (2020) State changes of the HORMA protein ASY1 are mediated by an interplay between its closure motif and PCH2. Nucleic Acids Res 48: 11521–11535

Ye Q, Kim DH, Dereli I, Rosenberg SC, Hagemann G, Herzog F, Toth A, Cleveland DW, Corbett KD (2017) The AAA+ ATPase TRIP13 remodels HORMA domains through N-terminal engagement and unfolding. EMBO J 36: 2419–2434

Ye Q, Rosenberg SC, Moeller A, Speir JA, Su TY, Corbett KD (2015) TRIP13 is a protein remodeling AAA+ ATPase that catalyzes MAD2 conformation switching. Elife 4:e07367

Yost S, de Wolf B, Hanks S, Zachariou A, Marcozzi C, Clarke M, de Voer R, Etemad B, Uijttewaal E, Ramsay E et al (2017) Biallelic TRIP13 mutations predispose to Wilms tumor and chromosome missegregation. Nat Genet 49: 1148–1151

Zanders S, Alani E (2009) The pch2Delta mutation in baker’s yeast alters meiotic crossover levels and confers a defect in crossover interference. PLoS Genet 5: e1000571

Zeng X, Li K, Yuan R, Gao H, Luo J, Liu F, Wu Y, Wu G, Yan X (2017) Nuclear Envelope Associated Chromosome Dynamics during Meiotic Prophase I. Front Cell Dev Biol 5: 121

Zhang Z, Li B, Fu J, Li R, Diao F, Li C, Chen B, Du J, Zhou Z, Mu J et al (2020) Bi-allelic Missense Pathogenic Variants in TRIP13 Cause Female Infertility Characterized by Oocyte Maturation Arrest. Am J Hum Genet 107: 15–23

Zickler D, Kleckner N (2015) Recombination, Pairing, and Synapsis of Homologs during Meiosis. Cold Spring Harb Perspect Biol 7: a016626

## References

1. Smith, A.V. and Roeder, G.S. (1997) The yeast Red1 protein localizes to the cores of meiotic chromosomes. J Cell Biol, 136: 957–967.

2. Penedos A, Johnson AL, Strong E, Goldman AS, Carballo JA, Cha RS (2015) Essential and Checkpoint Functions of Budding Yeast ATM and ATR during Meiotic Prophase Are Facilitated by Differential Phosphorylation of a Meiotic Adaptor Protein, Hop1. PLoS ONE 10: e0134297.

3. Herruzo, E., Santos, B., Freire, R., Carballo, J.A. and San-Segundo, P.A. (2019) Characterization of Pch2 localization determinants reveals a nucleolar-independent role in the meiotic recombination checkpoint. Chromosoma, 128: 297–316.

4. Sym M, Engebrecht JA, Roeder GS (1993) ZIP1 is a synaptonemal complex protein required for meiotic chromosome synapsis. Cell 72:365–378

